# RNF13 mediates pH- and Ca^2+^-dependent regulation of lysosomal positioning

**DOI:** 10.1101/2025.03.19.644102

**Authors:** Lan Thi Trinh, Anh Van Vu, Seunghye Shin, Chanhaeng Lee, Sang-Hee Park, Eun-Bee Cho, Yunseok Heo, Hyun Jin Kim, Sungjoo Kim, Jong-Bok Yoon

**Author notes:** Corresponsences to Sungjoo Kim or Jong-Bok Yoon. These authors contributed equally to the work (Sungjoo Kim, Jong-Bok Yoon).

## Abstract

Environmental factors such as extracellular pH (pH_e_) and nutritional status influence lysosomal localization and autophagy. However, the mechanisms by which pH_e_, intracellular pH (pH_i_), and Ca^2+^ levels coordinate the bidirectional transport of lysosomes remain poorly understood.

In this study, we identify RNF13 as a critical regulator of lysosomal positioning through its ubiquitin-dependent degradation of ARL8B. RNF13 activity is modulated by both pH_i_ and Ca^2+^ levels. Specifically, we demonstrate that Ca^2+^-activated apoptosis-linked gene 2 (ALG-2) promotes retrograde lysosomal transport while simultaneously increasing pH_i_ and decreasing lysosomal pH (pH_lys_). Elevated pH_i_ deprotonates RNF13 at His332, enabling its interaction with Ca^2+^-bound ALG-2. This interaction activates RNF13, which inhibits ARL8B-mediated anterograde lysosomal transport. Furthermore, we show that alkaline pH_e_ elevates pH_lys_, activating the lysosomal Ca^2+^ channel TRPML3. This activation enhances RNF13 activity, driving lysosomes to adopt a perinuclear localization. Thus, conditions such as starvation or alkaline pH_e_, which induce ALG-2 activation and pH_i_ elevation, facilitate RNF13- mediated ARL8B degradation. In contrast, under acidic pH_i_ conditions, RNF13 activity remains suppressed regardless of ALG-2 activation, leading to increased ARL8B levels. Additionally, we provide evidence linking the loss of RNF13 activity to developmental and epileptic encephalopathy-73, a neurological disorder characterized by severe developmental symptoms.

These findings deepen our understanding of lysosomal positioning mechanisms, highlighting the interplay between lysosomal Ca^2+^ release and dynamic changes in cytoplasmic and lysosomal pH.

## Introduction

Lysosomes and late endosomes (collectively referred to as lysosomes) are central to the degradation of macromolecules from both extracellular and intracellular sources, delivered via endocytosis and the autophagic pathway, respectively. Beyond their degradative role, lysosomes serve as critical regulators of energy metabolism and nutrient homeostasis by acting as a signaling hub^1–3^. Lysosomes are highly heterogeneous, differing significantly in size, shape, luminal pH, and subcellular localization depending on the physiological context. Perinuclear lysosomes are typically larger, more acidic, and exhibit higher degradative activity, but they are less motile compared to their peripheral counterparts.

The regulated intracellular distribution of lysosomes is essential for numerous cellular processes, including cancer cell invasion, antigen presentation, plasma membrane repair, and autophagy^4–6^. Lysosomal trafficking is governed by microtubule-based transport, interactions with other organelles^7–9^, and the composition of phosphatidylinositol phosphates (PIPs) in lysosomal membranes^10–14^. Microtubule-associated motor proteins mediate the bidirectional transport of lysosomes: kinesin facilitates anterograde transport toward the cell periphery, while dynein drives retrograde transport toward the perinuclear region. The directional movement of lysosomes is likely influenced by the force balance between these opposing motor proteins^15,16^, which are recruited to lysosomes through interactions with small GTPases and their effectors. RAB7, for instance, promotes dynein-mediated transport to the perinuclear region by interacting with its effector, Rab-interacting lysosomal protein (RILP)^15,17^. Conversely, RAB7 association with FYVE and coiled-coil domain-containing autophagy adaptor (FYCO1) facilitates kinesin-driven anterograde transport toward the cell periphery. Additional Rab GTPases, such as Rab26, Rab34, and Rab36, also recruit RILP to mediate perinuclear lysosomal clustering^18–22^. ARL8B is a key GTPase regulating anterograde lysosomal transport and is critical for diverse physiological functions^23–28^. It directly interacts with kinesin-3 or indirectly with kinesin-1 via its effector SKIP (also known as PLEKHM2) to facilitate anterograde transport^29^. BLOC-1-related complex (BORC) regulates the association of ARL8B to lysosomal membranes^30^. Interestingly, ARL8B can also participate in retrograde transport by recruiting effectors such as RUN- and FYVE-domain-containing proteins (RUFY3)^31,32^ or DENND6A^33^. However, the mechanisms by which RAB7 and ARL8B GTPases, along with their multiple effectors and adaptors, coordinate directional movement of lysosomes in response to physiological stimuli remain poorly understood. The relative levels of RAB7 and ARL8B are thought to play a pivotal role in determining lysosomal distribution^34^. Peripheral lysosomes are enriched in ARL8B but display lower levels of RAB7, highlighting a complex interplay between these GTPases in lysosomal positioning^34^.

Environmental factors, such as pH_e_ and nutrient availability, play critical roles in regulating lysosomal localization. An alkaline pH_e_, which leads to a corresponding increase in pH_i_ ^35–37^, promotes the perinuclear positioning of lysosomes^38,39^ and stimulates autophagy^37^. In contrast, an acidic pH_e_ exerts the opposite effects. Nutrient deprivation induces the redistribution of lysosomes to the perinuclear region, where they fuse with autophagosomes to facilitate cargo degradation. Conversely, under nutrient replete conditions, a significant portion of lysosomes move to the cell periphery, which facilitates mTORC1 activation by placing mTORC1 on lysosomes closer to nutrient-mediated signaling inputs into the cell^40^. Multiple regulators are involved in controlling the starvation-induced clustering of lysosomes in the perinuclear region^14,34,38,41–45^. Notably, starvation triggers an increase in intracellular pH (pH_i_), which impairs the recruitment of ARL8B to lysosomes and enhances retrograde lysosomal trafficking^41^. However, the mechanisms driving starvation-induced intracellular alkalinization and the reduced association of ARL8B with lysosomes remain poorly understood.

Nutrient deprivation activates the lysosomal Ca^2+^ channel TRPML1, a pivotal regulator of autophagy and lysosomal biogenesis. ALG-2, an effector of TRPML1, facilitates dynein-dependent perinuclear lysosomal positioning through Ca^2+^-dependent recruitment^42^. Elevated intracellular Ca^2+^ levels activate Ca^2+^/calmodulin-dependent protein kinase kinase β (CaMKKβ), which enhances autophagosome formation^46^, and calcineurin, which induces the nuclear translocation of transcription factor EB (TFEB)^47^. TFEB, a master regulator of lysosome function and autophagy, drives the expression of numerous target genes, including TMEM55B, which encodes a protein that recruits the dynein adaptor JIP4 to mediate retrograde lysosome transport^43^. Despite these insights, it remains unclear whether, or how, starvation-induced elevation of pH_i_ and lysosomal Ca^2+^ release are mechanistically linked to drive perinuclear lysosomal positioning (Figure S1A).

RING finger protein 13 (RNF13) is a type I transmembrane protein predominantly localized in lysosomes and a member of the PA-TM-RING E3 ligase family. This family is characterized by a signal peptide, a protease-associated (PA) domain, a transmembrane (TM) domain, and a RING-finger domain^48^. RNF13 plays a role in various biological processes, including skeletal muscle growth, neuronal development, and tumorigenesis^49^. Notably, missense variants in *RNF13* are associated with developmental and epileptic encephalopathy-73 (DEE-73), a severe neurological disorder^50^. These variants have been linked to impaired lysosomal localization^51^. Despite these insights, the molecular mechanisms underlying DEE-73 remain poorly understood. In this study, we aim to investigate how RNF13 regulates lysosomal positioning in response to pH_e_ changes and nutrient deprivation, and to investigate the molecular basis of DEE-73.

## Result

### RNF13-promoted ARL8B degradation is required for starvation- or alkaline pH_e_-induced perinuclear lysosomal positioning

To identify RNF13-specific ubiquitinated substrates, we employed a proximity-dependent biotin labeling method that we had previously developed^52–55^. This approach utilizes a fusion protein comprising RNF13 and the biotin ligase BirA. The biotin ligase selectively biotinylates an acceptor peptide fused to ubiquitin (AP-Ub), enabling the transfer of biotinylated AP-Ub to substrates in close proximity (Figure S1B). Biotinylated proteins were purified using streptavidin and analyzed as described in the STAR Methods section, leading to the identification of ARL8B as a potential RNF13 substrate (Figure S1C).

To validate whether RNF13 ubiquitinates and destabilizes ARL8B, HeLa cells were transfected with RNF13-HA either alone or in combination with ARL8B-FLAG. RNF13-HA expression reduced the levels of both endogenous and exogenous ARL8B in a dose-dependent manner, whereas the ligase-dead mutant RNF13 C243S (RNF13 CS)-HA had no effect (Figure 1A, S1D). This reduction in ARL8B levels was further corroborated by indirect immunofluorescence analysis, which showed a decrease in ARL8B in RNF13-transfected cells (Figure S1E). Treatment with the proteasome inhibitor MG132 or the lysosomal inhibitor NH_4_Cl increased endogenous ARL8B protein levels and mitigated the RNF13- mediated reduction of overexpressed ARL8B (Figure 1B). These results indicate that RNF13 promotes ARL8B degradation through both proteasomal and lysosomal pathways.

**Figure 1.**
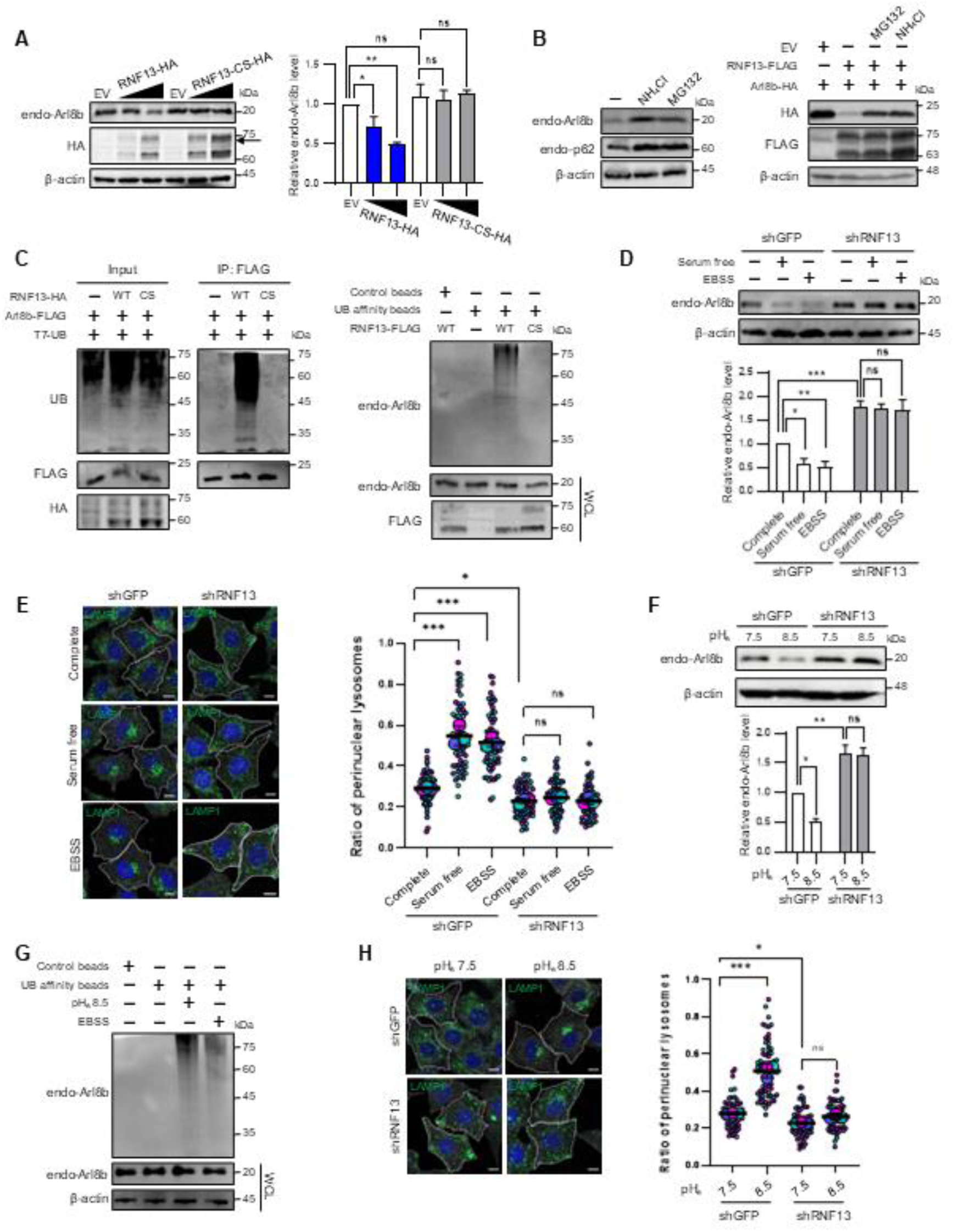
RNF13-promoted ARL8B degradation is required for starvation and alkaline pH_e_-induced perinuclear lysosomal positioning. (A) Western blot analysis of endogenous ARL8B in HeLa cells transfected with either an empty vector (EV) or plasmids encoding RNF13-HA or RNF13 C243S (CS)-HA at increasing concentrations. The arrow indicates the N-glycosylated form of RNF13. Protein levels were normalized to β-actin. Relative protein levels are shown as mean ± SD (n = 3). Statistical significance was assessed using one-way ANOVA with Dunnett’s multiple comparison test. (ns, not significant; *p < 0.05; **p < 0.01). (B) Western blot analysis of indicated proteins in HeLa cells treated with DMSO (-), MG132 (25 μM), or NH_4_Cl (20 mM) for 6 h of: (left panel) normal cells; (right panel) cells co-transfected with ARL8B- HA and either an EV or plasmid encoding RNF13-FLAG. Increase in p62 levels confirmed the effectiveness of MG132 and NH_4_Cl. (C) Western blot analysis for the ubiquitination assay of ARL8B. (Left panel) HeLa cells were transfected with plasmids encoding ARL8B-FLAG and T7-UB, along with either an EV, RNF13-HA, or RNF13 CS- HA plasmids. Cell lysates were subjected to immunoprecipitation (IP) using FLAG antibodies, followed by Western blotting. (Right panel) Lysates from cells transfected with EV, RNF13-FLAG, or RNF13 CS- FLAG were incubated with ubiquitin-affinity or control beads. Beads were collected, and the eluates were analyzed by Western blotting. For all ubiquitination assays, cells were pretreated with MG132 (25 μM) for 6 h before harvesting. Western blots were performed using the indicated antibodies. (D) Western blot analysis of endogenous ARL8B levels following starvation. Cell extracts from MCF7 cells stably expressing control shRNA (shGFP) or RNF13 shRNA (shRNF13) were immunoblotted after incubation with either complete, EBSS, or serum-free media for 2 h. Relative ARL8B levels, normalized to β-actin, were determined and presented as mean ± SD (n = 3). Statistical significance was assessed using two-way ANOVA with Tukey’s multiple comparisons test’s test. (ns, not significant; *p < 0.05; **p < 0.01; ***p < 0.001). (E) (Left panel) Immunofluorescence analysis of LAMP1 in the samples in (D) at 2 h. Samples were immunostained with LAMP1 for lysosomes, CD147 for cell boundary, and DAPI for nucleus. Scale bar, 10 μm. (Right panel) A SuperPlot shows the ratio of perinuclear LAMP1 to total LAMP1, determined through shell analysis (see the STAR Methods). For each condition, three independent experiments were performed. Over 20 cells were analyzed in individual experiment, which is distinguished by different colors. Small circles represent individual data points, while large circles denote the mean of each experiment. Horizontal lines represent the mean ± SD of the means from three independent experiments. Statistical significance was assessed by two-way ANOVA with Tukey’s test (ns, not significant; *p < 0.05; ***p < 0.001). (F) Western blot analysis of endogenous ARL8B in MCF7 cells stably expressing control shRNA (shGFP) or RNF13 shRNA (shRNF13) after incubation with alkaline media for 2 h. The relative ARL8B levels, normalized to β-actin, were determined and presented as mean ± SD (n=3). Statistical significance was analysed by two-way ANOVA with Tukey’s test (ns, not significant; *p < 0.05; **p < 0.01). (G) Western blot analysis for ubiquitination assay of endogenous ARL8B. HeLa cells were treated as indicated for 2 h and cell lysates incubated with either control or ubiquitin affinity beads as described in panel (C). The whole cell lysates and the eluates were analyzed by western blotting with indicated antibodies. (H) (Left panel) Immunofluorescence analysis of LAMP1 in the samples shown in (F) at 2 h. Samples were immunostained with LAMP1 for lysosomes, CD147 for cell boundary, and DAPI for nucleus. Scale bar, 10 μm. (Right panel) A SuperPlot illustrates the ratio of perinuclear LAMP1 to total LAMP1 as described in panel (E). Horizontal lines indicate the mean ± SD of the means from three independent experiments. Statistical significance was evaluated using two-way ANOVA with Tukey’s test (ns, not significant; *p < 0.05; ***p < 0.001).

RNF13-mediated ubiquitination of both endogenous and overexpressed ARL8B was confirmed through the detection of polyubiquitin chain formation. A characteristic ubiquitination smear was observed in the presence of wild-type RNF13 but not with the RNF13 CS mutant (Figure 1C). Furthermore, knockdown of RNF13 using RNF13-specific siRNAs resulted in an increase in endogenous ARL8B levels (Figure S1F), demonstrating that RNF13 negatively regulates ARL8B protein levels. This regulatory effect of RNF13 on ARL8B protein levels was also validated in MCF7 cells (Figure S1G). To further confirm the specificity of RNF13-mediated ubiquitination of ARL8B, we generated ARL8B mutants in which lysine residues at positions K131, K141, and K146 were substituted with arginine (K-to-R mutations). RNF13 failed to reduce the protein levels of the ARL8B K131/141/146R (3KR) mutant (Figure S1H) and was unable to ubiquitinate the mutant protein (Figure S1I). These findings identify K131, K141, and K146 as the primary ubiquitination sites on ARL8B. Collectively, these results demonstrate that RNF13 facilitates the ubiquitination of ARL8B, targeting it for degradation.

Overexpression of ARL8B promotes lysosomal localization toward the cellular periphery, while its knockdown induces lysosomal clustering in the perinuclear region^30^. Similarly, we observed that RNF13 overexpression causes perinuclear lysosomal clustering, whereas RNF13 knockdown results in lysosomal dispersal (Figure S1J-L). ARL8B knockdown has also been shown to enhance autophagosome-lysosome fusion and accelerate the degradation of autophagy substrates^41^. Consistent with this, depletion of RNF13 resulted in increased levels of p62 and LC3, indicative of impaired autophagic flux (Figure S2A). Conversely, RNF13 overexpression reduced their levels, suggesting enhanced autophagy (Figure S2B). Autophagosome-lysosome fusion assays further revealed that ARL8B overexpression and RNF13 knockdown inhibited autophagosome-lysosome fusion, whereas ARL8B knockdown and RNF13 overexpression promoted fusion (Figure S2C, D). These findings suggest that RNF13 regulates autophagy, likely by modulating ARL8B activity.

Changes in lysosomal positioning influenced by starvation and pH_i_ have been linked to alterations of ARL8B levels on lysosomes^42^. To elucidate the role of RNF13 in lysosomal positioning in response to nutrient and pH_e_, which affects pH_i_, we employed stable RNF13 knockdown in MCF7 cells using short hairpin RNA (shRNA). Incubation with Earle’s Balanced Salt Solution (EBSS) or serum-free media resulted in decreased ARL8B levels in control knockdown cells. However, this reduction was not observed in RNF13-knockdown cells during starvation (Figure 1D). Moreover, while starvation triggered retrograde movement of lysosomes toward perinuclear region in control knockdown cells, lysosomes remained dispersed throughout the cytoplasm in RNF13-knockdown cells (Figure 1E). Transient depletion of ARL8B via siRNA induced perinuclear lysosomal clustering in both control and RNF13-knockdown cells (Figure S3A, B). This suggests that lysosomal positioning alterations induced by RNF13 knockdown may result from corresponding increase in ARL8B levels.

To evaluate the effect of alkaline pH_e_ on lysosome positioning, we incubated cells at different pH_e_. We confirmed that changes in pH_e_ correspondingly induced alteration in pH_i_ (Figure S3C, D). In control knockdown cells, ARL8B levels were lower at pH_e_ 8.5 compared to pH_e_ 7.5, whereas this reduction was not observed in RNF13-knockdown cells (Figure 1F). MG132 or NH_4_Cl treatment inhibited RNF13-mediated degradation of endogenous ARL8B induced by alkaline pH_e_ and starvation (Figure S3E). The observed decrease in ARL8B levels correlated with increased ARL8B ubiquitination, which was enhanced by both alkaline pH_e_ and starvation (Figure 1G). Additionally, RNF13 depletion inhibited the perinuclear lysosomal clustering induced by alkaline pH_e_ (Figure 1H). Together, these findings suggest that RNF13 is essential for regulating perinuclear lysosomal positioning in response to starvation and alkaline pH_e_ by modulating ARL8B levels.

To investigate the kinetics of lysosome redistribution and time-dependent changes in ARL8B levels, we utilized time-lapse confocal microscopy. A stable cell line expressing LAMP1-GFP and ARL8B- mCherry was established for this purpose. Co-localization of LAMP1-GFP and ARL8B-mCherry during lysosomal transport throughout the cytoplasm confirmed their presence on lysosomes (Video S1, Figure S3F).

We evaluated whether LAMP1-GFP and ARL8B-mCherry mirrored the responses of endogenous LAMP1 and ARL8B to alkaline pH_e_ and starvation by assessing protein level changes via immunoblotting and analyzing lysosome distribution using fluorescence confocal microscopy 1-hour post treatment. Treatment with pH_e_ 8.5 or serum-free media induced perinuclear localization of both LAMP1-GFP and ARL8B-mCherry (Figure S3F). Additionally, the fluorescence intensity of ARL8B- mCherry decreased, while LAMP1-GFP signals remained unchanged (Figure S3F), a finding that was confirmed by immunoblotting (Figure S3G).

Time-lapse analysis revealed dynamic lysosomal motility and a gradual perinuclear redistribution during the 1-hour period following treatment with alkaline pH_e_ or starvation (Video S2, S3). Notably, in each cell analyzed, ARL8B-mCherry fluorescence steadily declined in response to these treatments (Figure S3H). In cells incubated with complete media at pH_e_ 7.5, ARL8B-mCherry fluorescence showed a decrease but to a lesser extent. This decrease in fluorescence is likely due to repeated laser exposure during the 1-hour imaging session, which differed from immunoblot analysis results (Figure S3G). Importantly, RNF13 depletion severely impaired alkaline pH_e_-induced lysosomal redistribution to the perinuclear region and decrease of ARL8B-mCherry fluorescence (Video S2, S4-6, and Figure S3I). These findings suggest that lysosome redistribution kinetics closely correlate with the observed decrease in ARL8B levels, supporting a link between lysosomal dynamics and ARL8B regulation.

We previously reported that over-expression and depletion of RNF167 affected ARL8B protein levels^54^. We therefore compared effects of RNF13 or RNF167 knockdown on regulation of lysosomal positioning and ARL8B levels in response to alkaline pH_e_. Unlike RNF13 depletion, RNF167 depletion did not severely inhibit alkaline pH_e_-induced perinuclear lysosomal positioning (Figure S3J). This observation was further validated by time-lapse analysis (Video S7 vs S5, S6). Alkaline pH_e_ led to a more pronounced decrease in ARL8B-mCherry fluorescence and protein levels in control and RNF167- depleted cells compared to RNF13-depleted cells (Figure S3K). Moreover, unlike RNF13, the RNF167- mediated reduction in ARL8B was not affected by acidic pH_e_ (Figure S3L). These findings indicate that RNF13 plays a pivotal role in regulating lysosomal positioning under physiological conditions that affect pH_i_. Based on these results, we focused on elucidating the mechanisms underlying RNF13 regulation.

### C-terminal tail of RNF13 regulates its activity toward ARL8B

Deletion mutants were used to identify the region(s) of RNF13 essential for ARL8B degradation (Figure 2A, B). While RNF13 (1-339) degraded ARL8B, comparable to full-length RNF13, RNF13 (1-318) and RNF13 (1-305) did not. Notably, RNF13 (1-286), which lacks the C-terminal region downstream of the RING domain, was still able to degrade ARL8B. In these C-terminal deletion mutants, the levels of the ligase-dead version with a C243S substitution were much higher than those without the substitution. MG132 increased the levels of these mutant proteins (Figure S4A), indicating that they possess RING-dependent auto-degradation activity, a typical feature of most E3 ubiquitin ligases, and that the C-terminal deletion did not cause significant folding defects. Furthermore, these data suggest that RNF13 (1-286) was sufficient for ARL8B recruitment and ubiquitination and that the C-terminal region, predicted to be unstructured by AlphaFold^56^, likely plays an active role in regulation of RNF13- mediated ARL8B degradation. Deletion analysis also revealed that while the region of RNF13 between residues 319 and 339 referred to as the PRR (Positive Regulatory Region), positively influenced ARL8B degradation, the region between residues 287 and 305, referred to as the NRR (Negative Regulatory Region), had the opposite effect. Additionally, internal deletion mutants RNF13 (Δ205-237) and RNF13 (Δ220-233) failed to show either auto-degradation or ARL8B degradation, while RNF13 (Δ205-219) and RNF13 (1-286, Δ205-219) mutants retained RING activity but were unable to degrade ARL8B (Figure 2B). Notably, a mutant with R to A substitutions at residues 215 and 217 exhibited impaired ARL8B degradation and interaction (Figures 2C, D). This suggests that these residues play a critical role in the recognition of ARL8B by RNF13.

**Figure 2.**
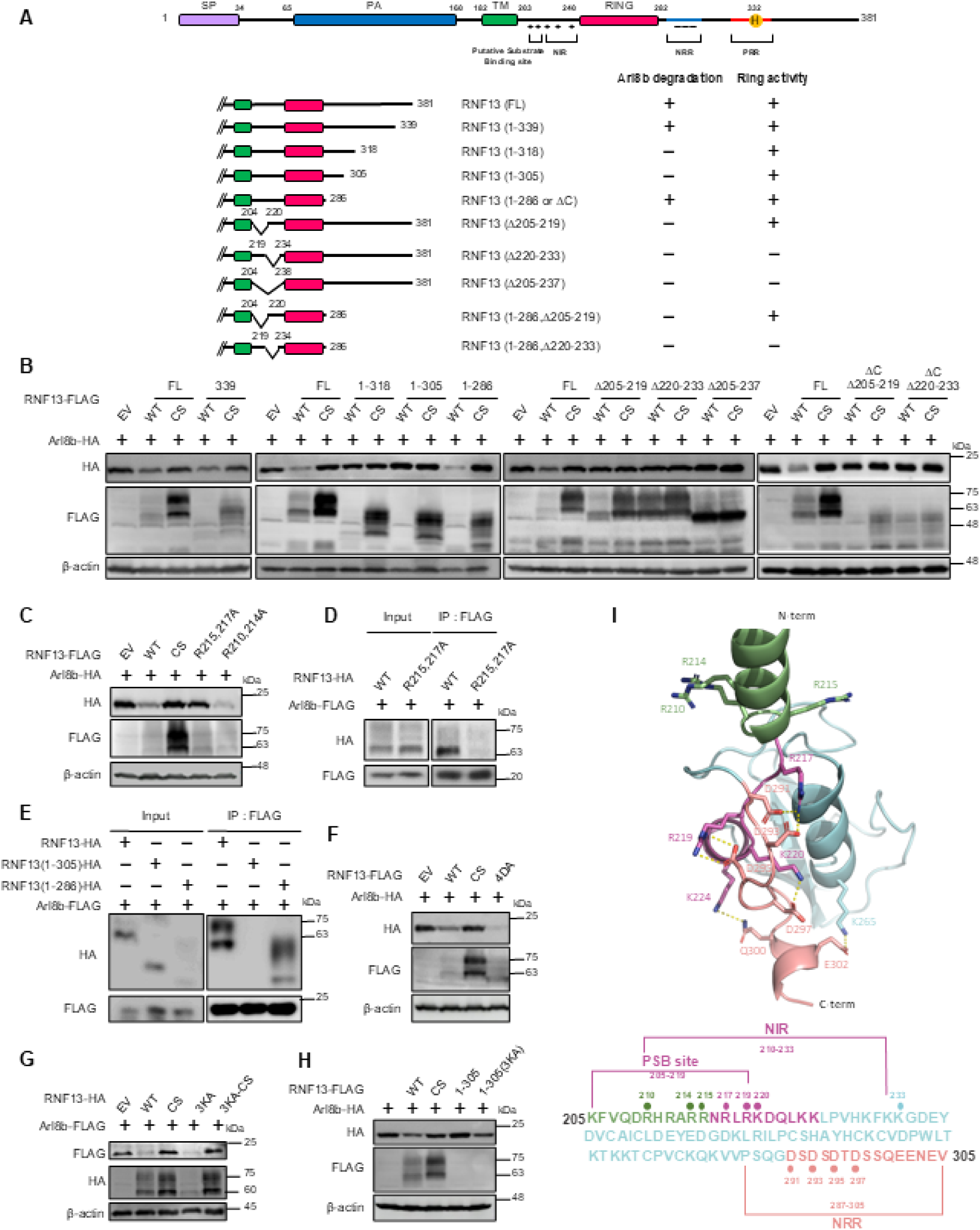
C-terminal tail of RNF13 regulates its activity toward ARL8B. (A) Schematic diagram of the RNF13 constructs with functional domains and their activities of ARL8B degradation and RING ligase. SP; signal peptide, PA; protease-associated domain, TM; transmembrane domain, RING; a really interesting new gene domain, NRR; negative regulatory region, NIR; NRR- interacting region, PRR; positive regulatory region. His332, the pH sensor of RNF13, is shown. (B-H) Western blot analysis of the indicated proteins in: (B, C) HeLa cells co-transfected with the plasmid encoding ARL8B-HA, along with the indicated RNF13 constructs for 24 h; (D, E) HeLa cells co- transfected with ARL8B-FLAG, along with the indicated RNF13 constructs for 24 h, treated with MG132 (25 μM) for 6 h prior to harvesting, and analyzed by IP with FLAG ; (F, G) HeLa cells overexpressing ARL8B with the indicated RNF13 constructs; (H) HeLa cells co-transfected with ARL8B-HA, along with indicated RNF13 constructs for 24 h. EV, empty vector; WT, construct with WT sequences; CS, construct with C243S mutation; R215,217A, construct with R-to-A substitution at 215 and 217; R210,214A, construct with R-to-A substitution at 210 and 214; 4DA, construct with D to A substitution at 291, 293, 295, and 297; 3KA, construct with K-to-A substitution at 220, 224, and 233; 3KA-CS, construct with 3KA and CS mutations; 1-305, a deletion construct containing AA 1 – 305; 1-305 (3KA), a deletion construct containing AA 1-305 with 3KA substitution . (I) Predicted structure of RNF13. The predicted RNF13 (aa. 205–305) model is depicted as a ribbon diagram. The N-terminal helical region (aa. 205–215) is colored in green, the basic-rich region (aa. 216– 225) in magenta, the acidic-rich region (aa. 291–305) in salmon, and the other residues are depicted in cyan. The side-chains of the residues involved in the interaction are represented as sticks, with CPK colors used for N and O atoms. Additionally, the side-chains of the residues R210, R214, and R215, used in the mutation study in panel C are also presented as sticks. A hydrogen bond is formed between K224 and Q300 (3.7 Å), while the other interactions are salt bridges. Salt bridges are observed between R217 and D291 (2.7 Å), R217 and D293 (2.8 Å), R219 and D295 (3.2–3.3 Å), K220 and D297 (2.7 Å), and K265 and E302 (2.9 Å). Each interaction is indicated by a yellow dotted line. The sequence of RNF13 (aa. 205–305) is shown below the structure, with the domains and key residues marked. The PSB site (aa. 205–219), NIR (aa. 210–233), and NRR (aa. 287–305) are indicated, with the critical residues involved in interactions highlighted as dots.

While the full-length RNF13 or RNF13 (1-286) lacking NRR was capable of ARL8B binding, NRR- containing RNF13 (1-305) failed to interact with ARL8B-HA (Figure 2E). NRR contains an unusually high number of negatively charged amino acids (7 of 19 residues with no positively charged ones). The role of negatively charged amino acids in NRR was investigated using a substitution mutant with A at positions 291, 293, 295, and 297 instead of D (named 4DA), which demonstrated increased ARL8B degradation activity (Figure 2F). Additionally, the region between the transmembrane and RING domains has a 24-amino acid stretch between residues 210 and 233 that contains 12 positively and 1 negatively charged residue. A mutant with K to A substitutions at positions 220, 224, and 233 (named RNF13 3KA) and RNF13 (1-305) 3KA increased ARL8B degradation (Figure 2G, H). Similar to wild-type RNF13 and the RNF13 CS mutant, the RNF13 CS 3KA mutant also interacted with ARL8B (Figure S4B), indicating that, unlike residues R215 and R217, the residues K220, K224, and K233 may not play a role in ARL8B interaction. These data suggest that intramolecular ionic interactions between the NRR and this positively charged region, referred to as the NIR (NRR-Interacting Region), may inhibit ARL8B binding by obstructing the overlapping ARL8B-binding site.

Given that the X-ray crystal structure of RNF13 spanning residues from 216 to 290 is available and the AlphaFold^56^ model predicts the region from 182 to 224 with high confidence, we utilized GalaxyWEB (https://galaxy.seoklab.org/) to dock the region from 291 to 305 into the combined RNF13 structure, integrating both the crystal and AlphaFold models. This approach allowed us to assess the plausibility of intramolecular interactions involving the NRR. The predicted structure suggests that the NRR interacts with a basic region that overlaps with the ARL8B-binding site (Figure 2I). These interactions may be critical for RNF13 regulation under conditions such as starvation and elevated pH_i_.

### Histidine 332 acts as a pH sensor to regulate the pH-dependent activity of RNF13

Given that cytosolic alkalinization decreased ARL8B levels in an RNF13-dependent manner, we investigated whether RNF13 activity acting on ARL8B is regulated by cytosolic pH. Epitope-tagged RNF13 and ARL8B were co-expressed in HeLa cells cultured in media at pH 7.4. Six hours post-transfection, the cells were transferred to fresh media at varying pH and incubated for an additional 18 hours. We observed that ARL8B-HA protein levels decreased with increasing pH_e_ when co-transfected with wild-type RNF13-FLAG (Figure 3A). In contrast, ARL8B-HA levels were only minimally affected by pH_e_ when co-transfected with the ligase-dead RNF13 mutant (RNF13 CS) or the empty vector. These results support the pH-sensitive activity of RNF13 toward ARL8B. Further analysis of RNF13 (1-286) demonstrated that this truncated form is not sensitive to changes in pH_e_ and exhibits greater activity than wild-type RNF13 at pH 6.5 (Figure 3A). As anticipated, RNF13 (1-286), but not full-length RNF13, induced perinuclear clustering of lysosomes under acidic pH_e_ conditions (Figure 3B). Additionally, the RNF13 3KA and RNF13 4DA mutants, where the interaction between the NRR and the NIR is likely disrupted, were pH-independent and showed higher activity compared to wild-type RNF13 (Figure 3C). Collectively, these findings suggest that the C-terminal region of RNF13 is integral to its pH-dependent regulation, with NRR-mediated negative regulation playing a pivotal role.

**Figure 3.**
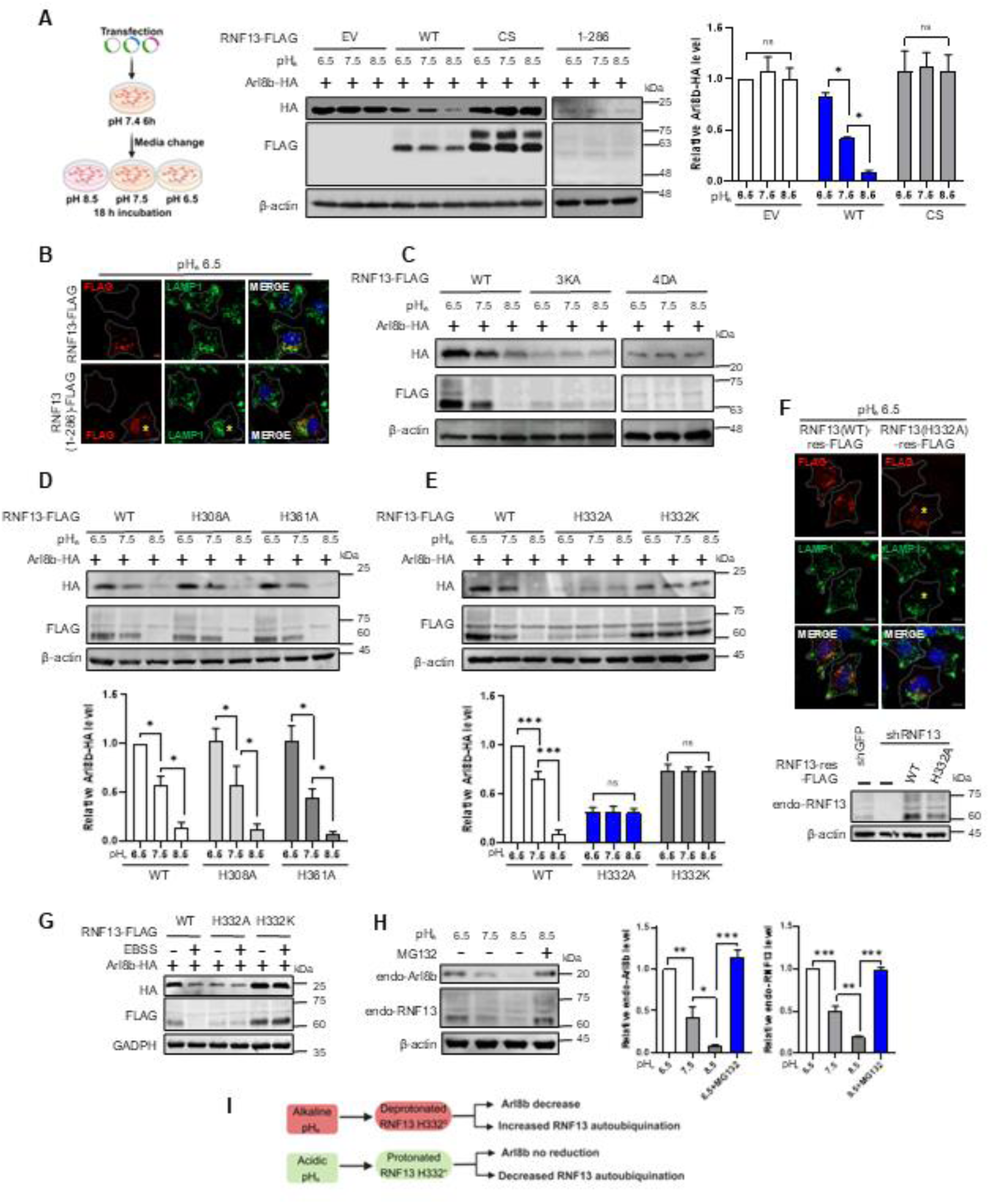
Histidine 332 acts as a pH sensor to regulate the pH-dependent activity of RNF13. (A) A Schematic of the experimental protocol (left panel) and Western blot analysis of lysates from HeLa cell co-transfected with plasmids expressing ARL8B-HA and either an empty vector (EV), RNF13- FLAG, RNF13 CS-FLAG, or RNF13 1-286-FLAG. Six hours post-transfection, the cells were incubated in fresh media at pH 6.5, pH 7.5, or pH 8.5 for 18 h. Cell lysates were subsequently analyzed by Western blot (middle panel). Relative ARL8B-HA levels, normalized to β-actin, were determined and presented as mean ± SD (n = 3) (right panel). Statistical significance: two-way ANOVA with Tukey’s test (ns, not significant; *p < 0.05). (B) Immunofluorescence analysis of indicated proteins in HeLa cells transfected with either RNF13- FLAG, or RNF13 1-286-FLAG as described in (A). Yellow asterisks indicate transfected cells with perinuclear lysosome positioning. Scale bar, 10 μm. (C) Western blot analysis of the indicated proteins in HeLa cells transfected with the indicated constructs for 6 h. After washing with PBS, the cells were incubated in fresh media at different pH for 18 h and then analyzed. (D, E) Western blot analysis and quantification of ARL8B. ARL8B-HA levels were normalized to β-actin. The relative ARL8B-HA levels were determined and presented as mean ± SD (n=3). Statistical significance was analyzed by two-way ANOVA with Tukey’s test (ns, not significant; *p < 0.05; ***p < 0.001). (F) Immunofluorescence of indicated proteins (upper panel) and Western blot analysis (lower panel) of HeLa cells stably expressing RNF13 shRNA (shRNF13) which were transfected with plasmids encoding the shRNA-resistant form of either the wild-type RNF13 or the RNF13 (H332A) mutant. Six hours later, cells were incubated in fresh media at pH 6.5 for 18 h and then analyzed as indicated. Scale bar, 10 μm. (G) Western blot analysis of the indicated proteins in HeLa cells co-transfected with plasmids expressing ARL8B-HA, in combination with either the wild-type RNF13 or the indicated mutants, followed by a 2- h incubation with EBSS. (H) Western blot analysis and quantification of endogenous ARL8B and RNF13 proteins in HeLa cells cultured in fresh media at different pH for 18 h. Cells were either untreated or treated with MG132 (25 μM) for 6 h prior to harvesting. Protein levels were normalized to β-actin, and relative levels are presented as mean ± SD (n = 3). Statistical significance was evaluated using two-way ANOVA with Tukey’s test. (*p < 0.05, **p < 0.01, ***p < 0.001). (I) Summary describing the role of H332 as a pH sensor in regulation of pH dependent RNF13 activity. The image was created using BioRender.com.

Histidine residues, with a near-neutral pKa, can be either positively charged or neutral at physiological pH, allowing them to sense pH changes. To investigate the roles of the three histidine (H) residues at positions 308, 332, and 361 in the C-terminal region of RNF13, substitution mutations with A or K were introduced at these positions. The mutants RNF13 H308A and RNF13 H361A demonstrated pH-dependent regulation similar to the wild-type RNF13 (Figure 3D) in that alkaline pH_e_-induced degradation of the substrate ARL8B and the enzyme itself. In contrast, RNF13 H332A and RNF13 H332K were not pH-dependent (Figure 3E). The pH-insensitive RNF13 H332A, but not wild-type RNF13, induced perinuclear lysosomal clustering at pH_e_ 6.5 (Figure 3F), suggesting that the H332A variant retains activity under acidic pH conditions. Given that starvation induces alkalization of pH_i_, the levels of co-expressed RNF13 and ARL8B decreased when cells were incubated in EBSS. In contrast, RNF13 H332A and RNF13 H332K mutants lost responsiveness to EBSS, as they did to pH changes (Figure 3G). To further investigate, we examined the effects of pH on endogenous RNF13 and ARL8B protein levels. Both RNF13 and ARL8B protein levels decreased as the pH increased (Figure 3H). Collectively, these findings suggest that H332 within the PRR may act as a pH sensor. Its deprotonation appears to activate RNF13, leading to increased auto-degradation of RNF13 and RNF13- dependent degradation of ARL8B (Figure 3I). The presence of pH-sensitive His332 in the PRR hinted that its protonated state may be closely associated with PRR function either directly by affecting intramolecular ionic interactions or indirectly through the recruitment of a regulatory protein that disrupts the NRR-NIR interaction.

### RNF13 is essential for inducing retrograde lysosomal transport by a TRPML agonist

In addition to its effect on pH_i_, starvation has been shown to activate the lysosomal Ca^2+^ channel TRPML1. This activation recruits the Ca^2+^-binding protein ALG-2 to lysosomes, where ALG-2 interacts with the dynactin-dynein complex, facilitating the retrograde transport of lysosomes^42^. To examine the role of RNF13 in TRPML-dependent lysosomal transport, we treated control and RNF13-knockdown cells with the TRPML agonist ML-SA1. In control cells, ML-SA1 reduced ARL8B protein levels, whereas this effect was absent in RNF13-knockdown cells (Figure 4A). Additionally, ML-SA1 induced perinuclear lysosomal accumulation in control cells (Figures S3F, G, Video S8) but not in RNF13- knockdown cells (Figure 4B). These findings suggest that RNF13-mediated degradation of ARL8B is required for ML-SA1-driven retrograde lysosomal trafficking.

**Figure 4.**
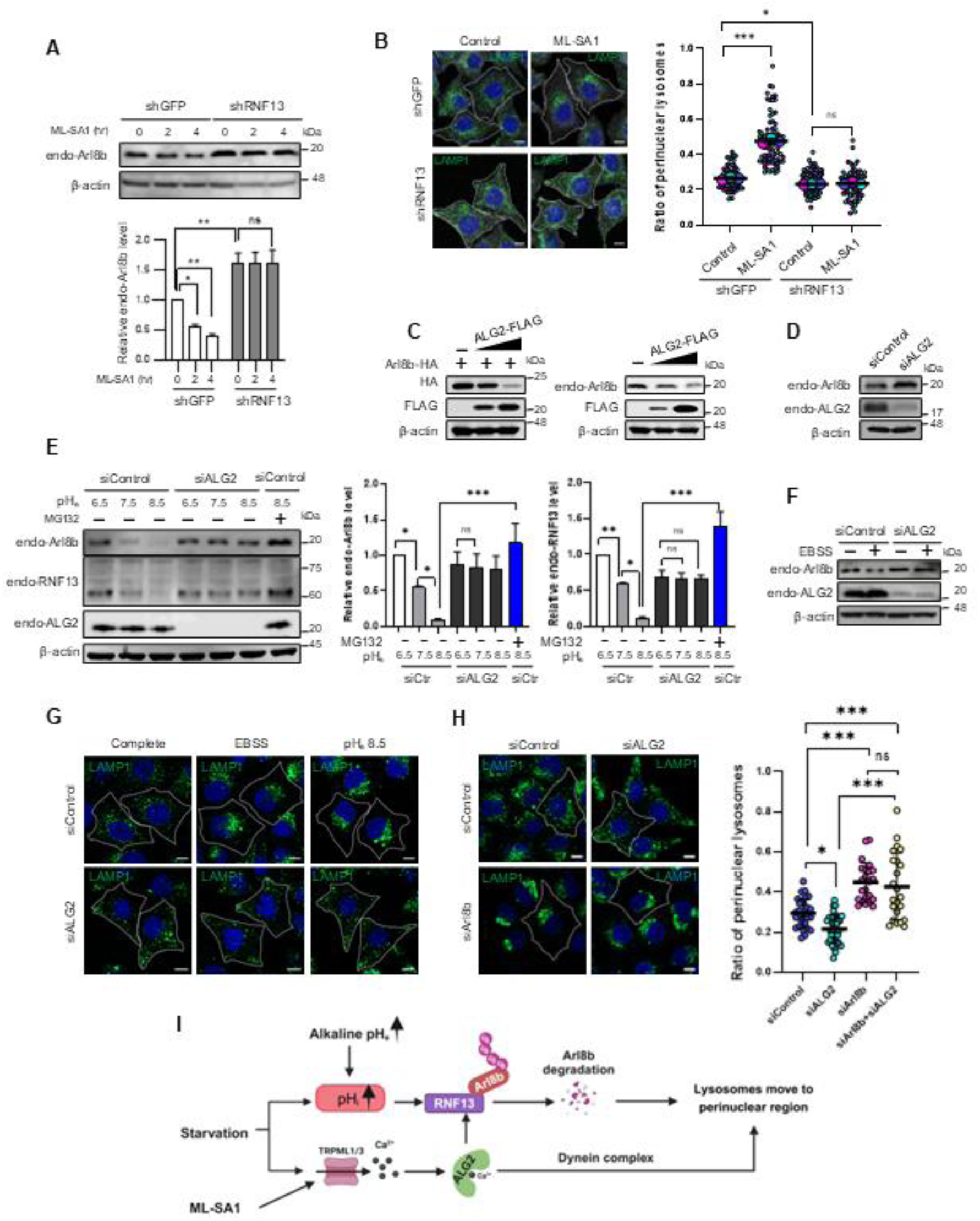
RNF13 is essential for inducing retrograde lysosomal transport by a TRPML agonist. (A) Western blot analysis and quantification of endogenous ARL8B protein levels in MCF7 cells stably expressing either control shRNA (shGFP) or RNF13 shRNA (shRNF13) treated with ML-SA1 (25 μM) for 0, 2, or 4 h. Endogenous ARL8B protein levels were normalized to β-actin, and relative levels are presented as mean ± SD (n = 3). Statistical significance: two-way ANOVA with Tukey’s test. (ns, not significant; *p < 0.05; **p < 0.01). (B) Immunofluorescence analysis of LAMP1 in the samples shown in (A) at 2 h (left panel). Scale bar, 10 μm. A SuperPlot depicting the ratio of perinuclear LAMP1 to total LAMP1, quantified using shell analysis of the immunofluorescence signals as described in Figure 1E (right panel). Horizontal lines represent mean ± SD (n = 3). Statistical significance was assessed by two-way ANOVA with Tukey’s test (ns, not significant; **p < 0.01; ***p < 0.001). (C) Western blot analysis of indicated proteins in HeLa cells transfected with increasing amounts of plasmid encoding FLAG-ALG-2, either in combination with a ARL8B-HA plasmid (left panel) or alone (right panel). The blots were probed with the indicated antibodies. (D) Western blot analysis of endogenous ARL8B and ALG-2 in HeLa cells transfected with either control siRNA or ALG-2-specific siRNA for 72 h and then subjected to immunoblotting. (E) Western blot analysis and quantification of the indicated proteins in HeLa cells transfected with either control siRNA or ALG-2-specific siRNA for 72 h and subsequently incubated in media at pH 6.5, pH 7.5, or pH 8.5 for 18 h. For comparison, MG132 (25 μM) was added to siControl cells six hours before collection. Protein levels were normalized to β-actin and relative endogenous protein levels were quantified from three independent Western blots (n = 3). Bars represent the mean ± SD. Statistical significance two-way ANOVA with Tukey’s test. (ns, not significant; *p < 0.05; **p < 0.01; ***p < 0.001). (F) Western blot analysis of endogenous ARL8B and ALG-2 in HeLa cells transfected with either control siRNA or ALG-2-specific siRNA for 72 h, and then followed by a 2 h-incubation in complete media, or EBSS. (G) Immunofluorescence analysis of LAMP1 in HeLa cells to assess the effects of ALG-2 knockdown on lysosome positioning under starvation (EBSS) or alkaline pH_e_ conditions. (H) Immunofluorescence analysis of LAMP1 in HeLa cells transfected with either control siRNA, ALG-2-specific siRNA (siALG-2), ARL8B-specific siRNA (siARL8B), or a combination of siALG-2 and siARL8B for 72 h (left panel). The ratio of perinuclear LAMP1 to total LAMP1 was quantified through shell analysis (right panel). Lysosome distribution was analyzed in over 20 cells per condition. Data are presented as mean ± SD, determined by one-way ANOVA with Tukey’s test. (ns, not significant; *p < 0.05; ***p < 0.001). (I) Summary describing that increased RNF13-mediated ARL8B degradation facilitates the perinuclear accumulation of lysosomes in response to starvation and ML-SA1 treatment. The image was created using BioRender.com.

Given the reduction in RNF13 activity at acidic pH_i_, we investigated whether ML-SA1 could induce ARL8B reduction and retrograde lysosomal transport under acidic pH_e_ 6.5. ML-SA1 promoted TFEB nuclear translocation at pH_e_ 6.5, confirming TRPML1 activation (Figure S4C). However, ML-SA1 treatment at pH_e_ 6.5 failed to induce perinuclear lysosomal clustering (Figure S4D). Additionally, while ML-SA1 reduced ARL8B levels at pH_e_ 7.5, this effect was absent under acidic pH_e_ conditions (Figure S4E). These findings suggest that TRPML1 activation alone is insufficient to drive retrograde lysosomal transport at acidic pH_i_, likely due to impaired RNF13-mediated ARL8B degradation. We confirmed that the TRPML antagonist ML-SI3 suppressed perinuclear lysosomal clustering under conditions of serum-free media or stimulation with ML-SA1, and found that ML-SI3 inhibited alkaline pH_e_-induced lysosomal redistribution to the perinuclear region (Figure S4F). Concurrently, ML-SI3 treatment blocked the reduction in ARL8B levels by these treatments (Figure S4G). These findings suggest that lysosomal Ca^2+^ release is crucial for RNF13 activation in response to alkaline pH_e_ or starvation.

Previous studies showed that ALG-2 overexpression led to the perinuclear positioning of lysosomes, whereas ALG-2 knockout resulted in their peripheral distribution^42^. We thus examined whether ALG-2 affects ARL8B protein levels. ALG-2 overexpression decreased both exogenous and endogenous ARL8B levels (Figure 4C), while ALG-2 knockdown increased endogenous ARL8B levels (Figure 4D). Importantly, ALG-2 knockdown prevented the reduction of endogenous RNF13 and ARL8B protein levels induced by alkaline pH_e_ (Figure 4E). Similarly, ALG-2 knockdown inhibited the decrease in ARL8B levels induced by starvation (Figure 4F). Furthermore, ALG-2 depletion blocked perinuclear lysosomal clustering induced by either starvation or alkaline pH_e_ (Figure 4G). To determine whether the peripheral lysosomal positioning observed with ALG-2 depletion was due to increased ARL8B levels, we simultaneously depleted both proteins. Perinuclear lysosomal clustering was observed in cells depleted of both ARL8B and ALG-2, suggesting that the increase in ARL8B levels drives peripheral lysosome positioning (Figure 4H). Collectively, these findings indicate that RNF13-mediated degradation of ARL8B and perinuclear lysosomal positioning are dependent on lysosomal Ca^2+^ release and the Ca^2+^-sensitive adaptor ALG-2 (Figure 4I).

### Starvation induces cytoplasmic alkalinization in a TRPML- and ALG-2-dependent manner

Since pH-sensitive RNF13 is essential for ML-SA1-induced retrograde lysosomal movement, we investigated the effect of ML-SA1 on pH_i_, and subsequently on RNF13 activity. The pH_i_ was monitored using a ratiometric pH probe consisting of the pH-sensitive GFP variant, Super-Ecliptic pHluorin, fused to the pH-insensitive mCherry^57,58^, which is well-suited for measuring relative pH changes (Figure S5A). Starvation induced a time-dependent increase in pH_i_, rising from 7.4 to 7.8 after 2-h EBSS incubation (Figure 5A). Remarkably, ML-SA1 caused cytosolic alkalinization to a comparable extent in cells cultured in complete media (Figure 5B). Inhibition of V-ATPase by bafilomycin A1 (BafA1) completely blocked EBSS- and ML-SA1-induced elevation of pH_i_ (Figure 5C), suggesting that V-ATPase is likely involved in the cytosolic alkalinization observed in response to starvation or TRPML activation. Since V-ATPase activity affects both cytoplasmic and lysosomal pH, we thus investigated the potential alteration of lysosomal pH (pH_lys_) by EBSS or ML-SA1 using pH_lys_ sensor cell line, which expresses a fusion protein of pH-sensitive monomeric teal fluorescent protein 1 (mTFP1) in the lumen of lysosomes and pH-insensitive mCherry on the cytosolic side^59^ (Figure S5B). Both EBSS and ML-SA1 reduced pH_lys_ from 4.7 to below 4.3 (Figure 5D), demonstrating that these treatments simultaneously induced a decrease in pH_lys_ and an increase in pH_i_. Additionally, ML-SI3 lowered pH_i_ from 7.4 to 7.2 in control cells, presumably by blocking a basal TRPML channel activity, and negated the effects of EBSS and ML-SA1 on pH_i_ by decreasing it to 7.2 (Figure 5E, S5C, S5D). Collectively, these results suggest that the cytosolic alkalinization and lysosomal acidification induced by starvation are primarily driven by TRPML channel activation.

**Figure 5.**
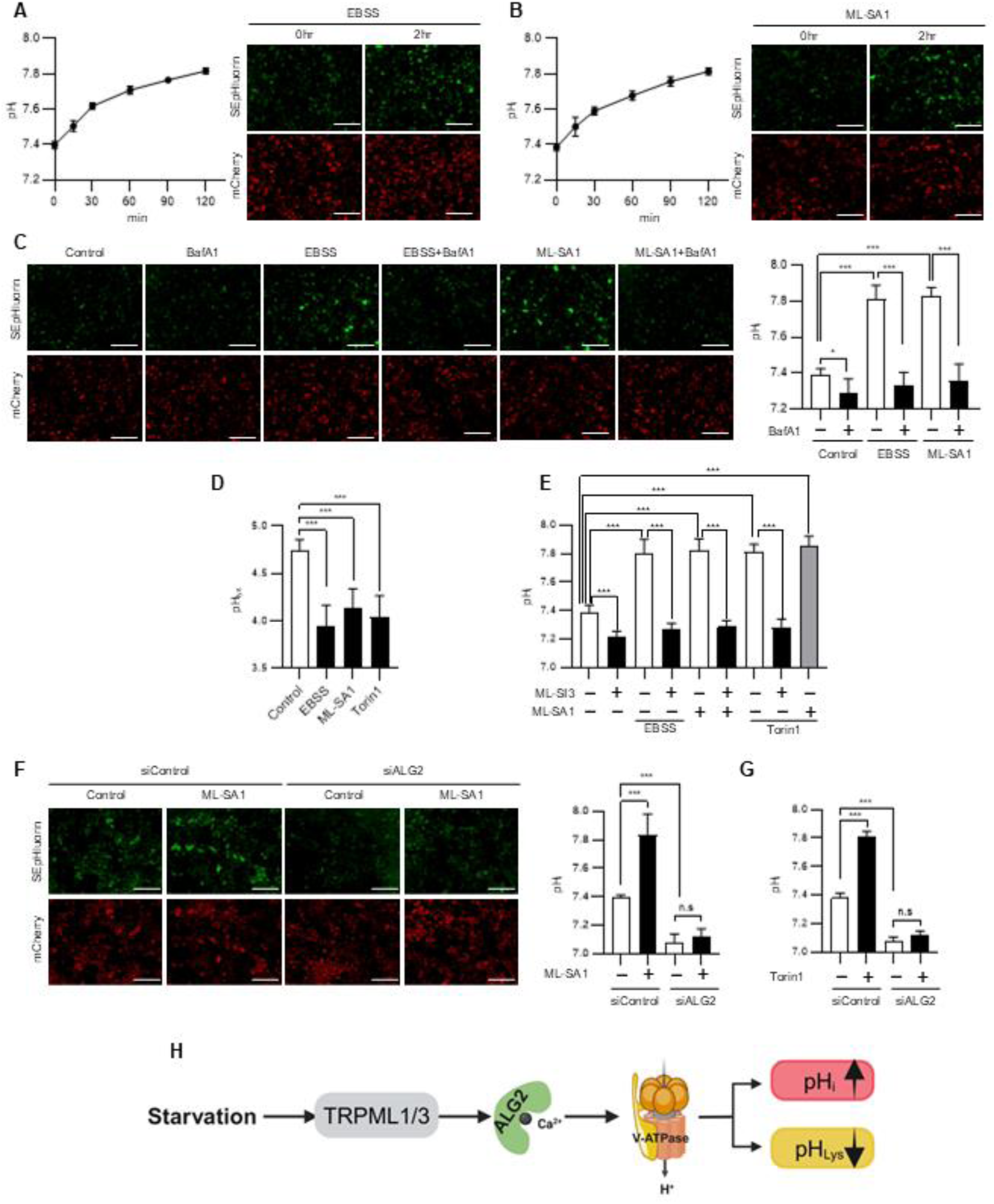
Starvation induces cytoplasmic alkalinization in a TRPML- and ALG-2-dependent manner. (A, B) pH_i_ changes in HeLa cells stably expressing the pH_i_ sensor mCherry-SEpHluorin (pH_i_ sensor cell line) were measured following a 2-hour treatment with EBSS (A) or ML-SA1 (25 μM) (B). SEpHluorin (green) and mCherry (red) signals were recorded, and the green-to-red ratio was converted to pH_i_ using a calibration curve (Figure S5A). The mean pH_i_ was calculated from three fields per condition, and these values were used to derive the mean pH_i_ ± SD based on three replicates. (C) pH_i_ measured in pH_i_ sensor cells after 2-hour incubation with DMSO, EBSS, or ML-SA1 (25 μM) following pre-incubation with DMSO (control) or BafA1 (10 nM) for 30 minutes. (Left panel) Immunofluorescence analysis. (Right panel) pH_i_ values were derived and calculated as described in (A). Statistical significance was assessed using two-way ANOVA with Tukey’s test (*p < 0.05; ***p < 0.001). (D) pH_lys_ measured in pH_lys_ sensor cells treated with DMSO (control), EBSS, ML-SA1 (25 μM), or Torin1 (400 nM) for 2 h. mTFP (green) and mCherry (red) signals from HeLa cells expressing mTFP1– hLAMP1–mCherry were converted to pH_lys_ using a calibration curve (Figure S5B). Mean pH_lys_ ± SD was calculated from three independent experiments, with over 20 cells observed per experiment. Statistical significance was determined by one-way ANOVA with Dunnett’s test. (∗∗∗p ≤ 0.001). (E) pH_i_ measured in pH_i_ sensor cells incubated in EBSS or treated with ML-SA1 (25 μM), or Torin1 (400 nM) for 2 h, following pretreatment with DMSO (control) or ML-SI3 (25 μm) for 30 min. SEpHluorin (green) and mCherry (red) signals were converted to pH_i_. The mean pH_i_ was calculated and statistical significance was assessed by two-way ANOVA with Tukey’s test (***p < 0.001). (F, G) pH_i_ measured in pH_i_ sensor cells transfected with control siRNA (siControl) or ALG-2 siRNA (siALG-2) for 72 h, followed by treatment with DMSO, or ML-SA1 (25 μM) (F) or Torin1 (400 nM) (G) for 2 h. The mean pH_i_ was calculated as described in (A), and statistical significance was assessed by two-way ANOVA with Tukey’s test. (ns, not significant; ***p < 0.001). (H) Summary of results in Figure 5. The image was created using BioRender.com.

A recent report showed that mTOR inhibition by Torin1 induced a decrease in pH_lys_ through upregulation of V-ATPase assembly and activity^60^. We confirmed this observation (Figure 5D), along with a simultaneous increase in pH_i_ (Figure 5E). This increase in pH_i_ was blocked by ML-SI3, suggesting that it was mediated by TRPML channel activation. Notably, co-treatment of Torin1 and ML-SA1 did not produce an additive or synergistic effect on pH_i_, indicating that both treatments likely act through a common target (Figure 5E).

We further investigated the role of ALG-2 in regulating pH_i_ and pH_lys_. Overexpression of wild-type ALG-2, but not a Ca^2+^ binding-deficient ALG-2 mutant, resulted in cytosolic alkalinization and lysosomal acidification (Figure S5E, F). Additionally, ALG-2 knockdown inhibited ML-SA1 induced pH_i_ increase (Figure 5F), indicating that Ca^2+^-activated ALG-2 is essential for TRPML channel- mediated cytosolic alkalinization. Similarly, ALG-2 knockdown prevented Torin1-induced cytosolic alkalinization (Figure 5G). Overall, these findings suggest that starvation triggers TRPML channel- dependent activation of ALG-2, which enhances V-ATPase activity, leading to a simultaneous decrease in pH_lys_ and increase in pH_i_ (Figure 5H).

### Ca^2+^-bound ALG-2 activates RNF13 via a His332 deprotonation-dependent interaction with PRR that relieves NRR-mediated repression

As Ca^2+^-activated ALG-2 was essential for RNF13 activation, we monitored relative cytosolic Ca^2+^ levels using a Ca^2+^ sensor GCaMP6f^61^. EBSS and ML-SA1 caused a continuous increase in GCaMP6f signals over a 2-h period (Figure 6A, B, S6A). We subsequently compared changes in cytosolic Ca^2+^ levels and pH_i_ at 2 h after treatment with ML-SA1, EBSS, serum-free media, or Torin1 (Figure 6C). Starvation and ML-SA1 increased GCaMP6f fluorescence ΔF/F0 to 0.44–0.63, whereas Torin1-induced a ΔF/F0 increase of no more than 0.11. Notably, all four conditions led to an increase in pH_i_ to 7.8. The observed Ca^2+^ increase indicates that Ca^2+^ release triggered by starvation or ML-SA1 treatment was significantly greater than that induced by mTOR inhibition. We further confirmed that a 2-h Torin1 treatment did not reduce ARL8B levels or significantly alter lysosomal positioning^41,45^ (Figure S6B). Additionally, RNF13 activation in Torin1-treated cells was evaluated by elevating cytosolic Ca^2+^ levels using thapsigargin, a SERCA inhibitor. Although thapsigargin effectively increased cytosolic Ca^2+^ levels, it did not produce a noticeable effect on pH_i_ (Figure S6C). Co-treatment with Torin1 and thapsigargin resulted in reduction in ARL8B protein levels and led lysosomes to the perinuclear region (Figure S6B), suggesting that cytosolic alkalinization induced by Torin1 requires elevated cytosolic Ca^2+^ levels, achieved using thapsigargin, for RNF13 activation. These effects were not visible with thapsigargin alone (Figure S6B).

**Figure 6.**
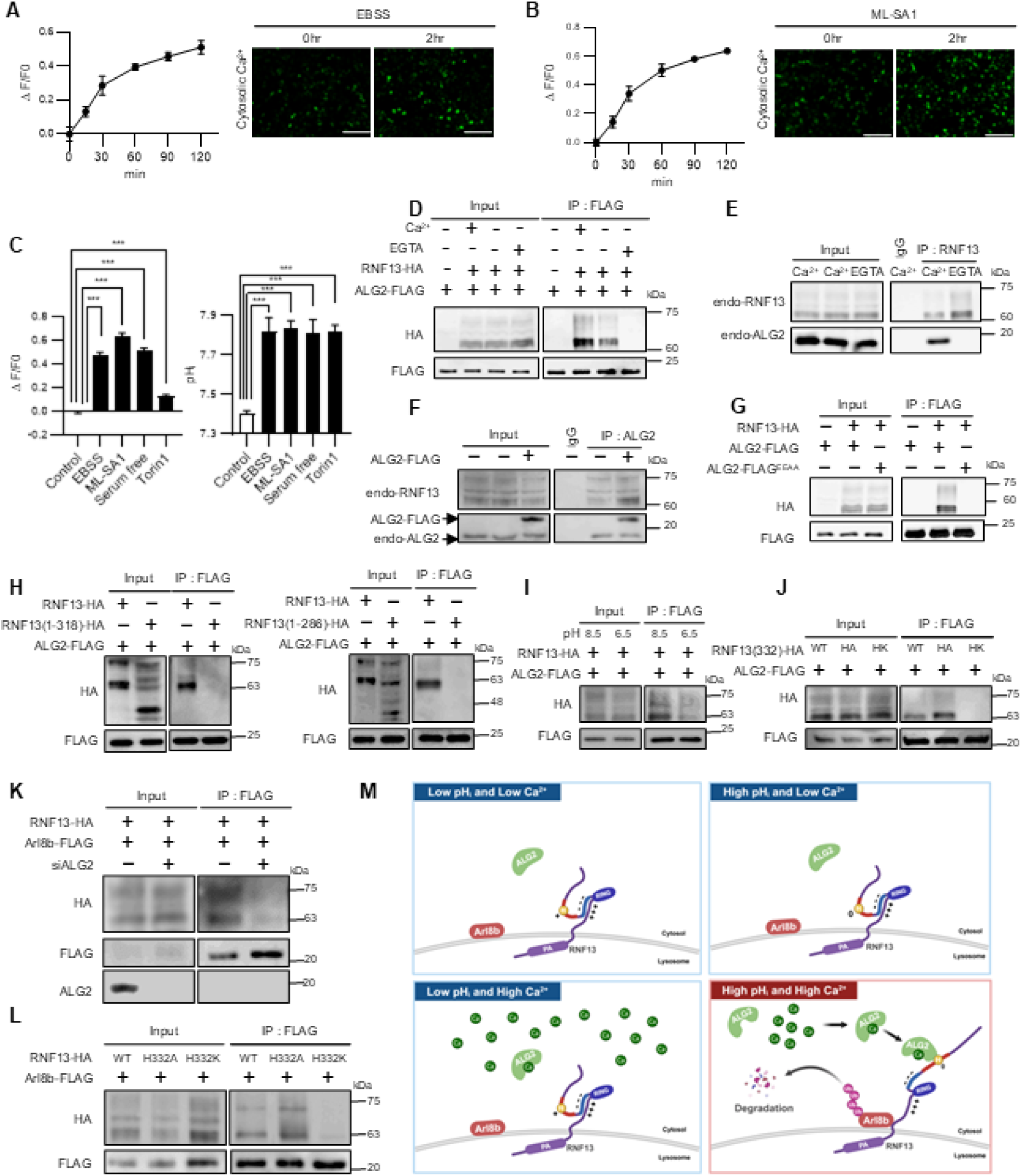
Ca^2+^-bound ALG-2 activates RNF13 via a His332 deprotonation-dependent interaction with PRR that relieves NRR-mediated repression. (A, B) Cytosolic Ca^2+^ levels measured following a 2-h treatment with EBSS (A) or ML-SA1 (25 μM) (B) using HeLa cells stably expressing the calcium sensor GCaMP6f (Ca^2+^ sensor cell line) as described in the STAR Methods. Data were collected from three independent experiments, each with three fields per measurement. Data were normalized to the initial time point for the same replicates. (C) Ca^2+^ levels in Ca^2+^ sensor cells (left panel) and pH_i_ in pH_i_ sensor cells (right panel) were analyzed after 2-hour incubation with DMSO (control), EBSS, ML-SA1 (25 μM), serum-free media, or Torin1 (400 nM). ΔF/F0 was measured in three fields per condition, and the mean ΔF/F0 ± SD was calculated from three replicates as described in the STAR Methods. Mean pH_i_ ± SD was determined as outlined in Figure 5A. Statistical significance was assessed using one-way ANOVA with Dunnett’s test (***p ≤ 0.001). (D-L) Co-Immunoprecipitation (Co-IP) analysis of the indicated proteins in lysates of HeLa cells: (D) co-transfected with the RNF13-HA and ALG-2-FLAG plasmids and analyzed by Co-IP with FLAG antibody in the absence or presence of 100 μm Ca^2+^ or 5 mM EGTA in the lysis buffer, (E) incubated with RNF13 antibodies and protein G-conjugated magnetic beads in the presence of 100 μm CaCl_2_ or 5 mM EGTA, (F) transfected with empty vector or ALG-2-FLAG and Co-IP with ALG-2 antibodies, (G) transfected with RNF13-HA, along with either ALG-2-FLAG or ALG-2^E47A/E114A^-FLAG (ALG-2^EEAA^) and Co-IP with FLAG antibody, (H) transfected with ALG-2-FLAG in combination with either RNF13 (1-318) or RNF13 (1-286) and Co-IP with FLAG antibody, (I) transfected with ALG-2-FLAG and RNF13-HA. Cell lysis and Co-IP with FLAG antibody were performed using buffer at pH 8.5 or pH 6.5, (J) co-transfected with ALG-2-FLAG, along with different RNF13(332) mutant constructs and Co-IP with FLAG antibody, (K) transfected with control siRNA(-) or ALG-2 siRNA (+) for 48 h, followed by transfection with RNF13-HA, and ARL8B-FLAG for 24 h, and Co-IP with FLAG antibody, (L) co- transfected with ARL8B-FLAG, along with the different RNF13 (332) constructs and Co-IP with FLAG antibody. (M) A proposed model for RNF13-mediated ARL8B degradation under conditions of different pH_i_ and Ca^2+^ levels. The image was created using BioRender.com.

We next performed co-immunoprecipitation (co-IP) assays to investigate the interaction between RNF13 and ALG-2 since RNF13 activation requires His332 deprotonation and Ca^2+^-bound ALG-2. Both overexpressed and endogenous RNF13 and ALG-2 were found to interact with each other in a Ca^2+^ dependent manner (Figure 6D, E). Overexpression of ALG-2 resulted in a greater amount of RNF13 being co-precipitated, suggesting an enhanced interaction under these conditions (Figure 6F). Furthermore, a Ca^2+^-binding-defective ALG-2 mutant (ALG-2-FLAG^EEAA^) was unable to bind to RNF13 (Figure 6G). Additionally, RNF13 variants lacking PRR, specifically RNF13 (1-318) and RNF13 (1-286), failed to interact with ALG-2, unlike the full-length RNF13 (Figure 6H). Notably, a greater amount of RNF13 was co-precipitated with ALG-2 at pH 8.5 compared to pH 6.5 (Figure 6I), suggesting that ALG-2 binding may require H332 deprotonation in the PRR. Indeed, while ALG-2 pulled down more RNF13 H332A mutant than the wild-type protein, it failed to interact with the RNF13 H332K mutant (Figure 6J). These findings indicate that H332 deprotonation in the PRR likely enhances the interaction between RNF13 and Ca^2+^-bound ALG-2.

We further explored the role of ALG-2 in mediating the interaction between RNF13 and ARL8B. Depletion of ALG-2 using siRNA significantly reduced the interaction between RNF13 and ARL8B (Figure 6K). Additionally, the interaction of the RNF13 H332A mutant with ARL8B was enhanced compared to wild-type RNF13, while the H332K mutant weakened the RNF13-ARL8B interaction (Figure 6L). These results suggest that the binding of ALG-2 to RNF13 facilitates the interaction of RNF13 with ARL8B.

Based on these findings, we propose that low cytosolic Ca^2+^ levels or enhanced protonation of H332 at acidic pH_i_ disrupts ALG-2 binding to the H332-containing PRR. This allows the NRR to intramolecularly interact with the NIR, which overlaps the ARL8B binding site, thereby inhibiting ARL8B interaction with RNF13. As a result, ARL8B stabilization occurs, promoting peripheral lysosomal positioning. Conversely, under conditions of high cytosolic Ca^2+^ levels and increased deprotonation of H332 at alkaline pH_i,_ ALG-2 binds to the PRR, preventing the NRR from engaging the NIR. This permits the interaction between RNF13 and ARL8B, facilitating RNF13-mediated ubiquitination and subsequent degradation of ARL8B, leading to perinuclear lysosomal positioning (Figure 6M). Given that RNF13 auto-degradation is influenced by pH_i_ and cytosolic Ca^2+^ levels, the same conformational changes in RNF13 may affect the binding of ubiquitin-loaded E2 enzymes.

### Alkaline pH_e_ increases cytosolic Ca^2+^ via TRPML3 activation

Since alkaline pH_e_ induces RNF13- and ALG-2-dependent perinuclear lysosomal positioning, we investigated whether alkaline pH_e_ also elevates cytosolic Ca^2+^ levels in addition to increasing pH_i_. Incubation at pH_e_ 8.5 raised pH_i_ to 7.8 and increased cytosolic Ca^2+^ with a ΔF/F_0_ of 0.5 (Figure 7A). ML-SI3 blocked the alkaline pH_e_-induced increase in cytosolic Ca^2+^ (Figure 7A). Additionally, ML-SI3 slightly reduced pH_i_ under pH_e_ 7.5 conditions, likely due to the inhibition of basal TRPML channel activity. These findings suggest that alkaline pH_e_ triggers lysosomal Ca^2+^ release through the activation of TRPML channels.

**Figure 7.**
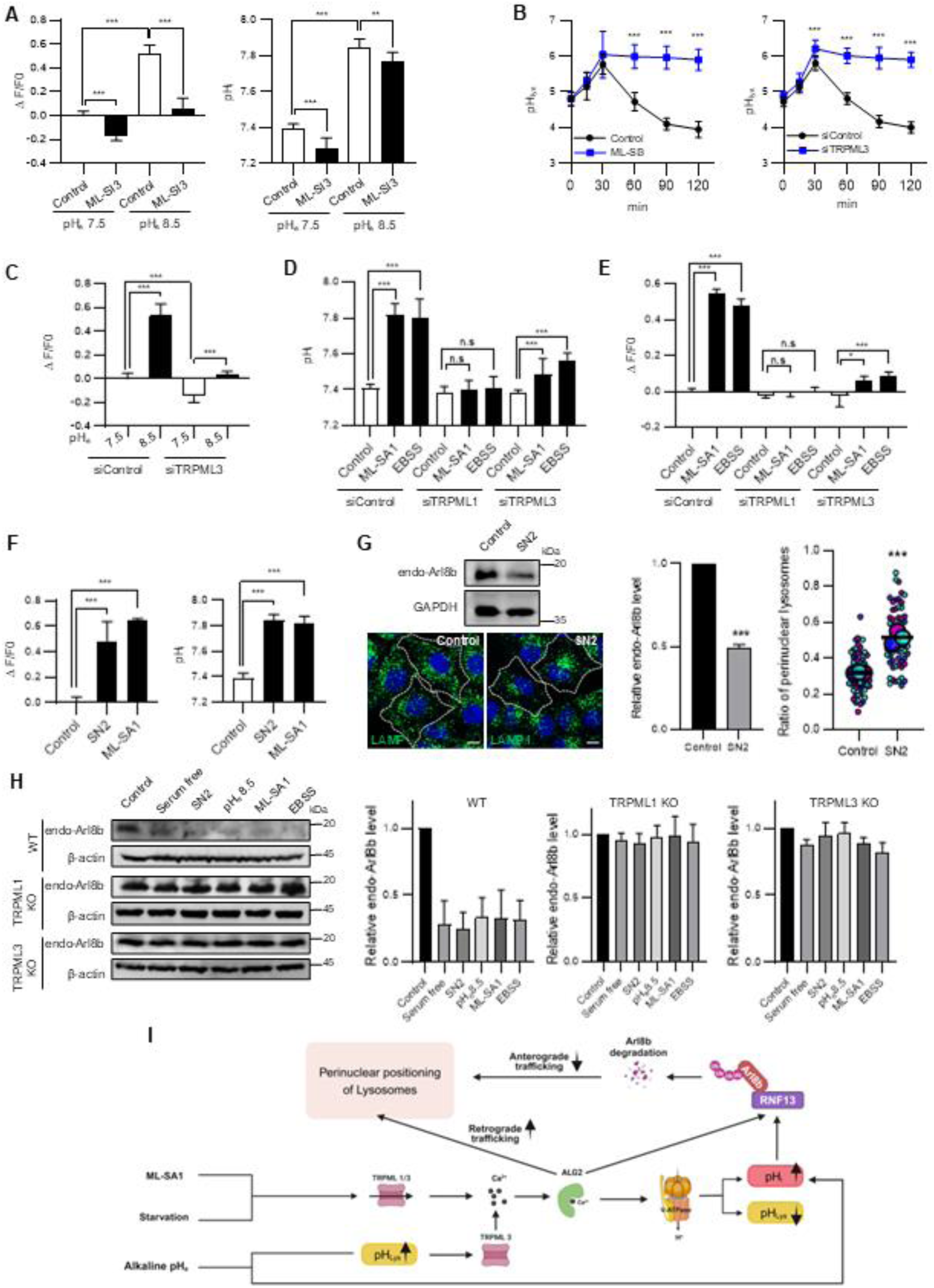
Alkaline pH_e_ increases cytosolic Ca^2+^ via TRPML3 activation. (A) Ca^2+^ levels in Ca^2+^ sensor cells (left panel) and pH_i_ in pH_i_ sensor cells (right panel) analyzed after a 2h incubation with media at different pH, following 30-min pretreatment with DMSO (control) or ML- SI3 (25 μM). ΔF/F0 was measured in three fields per condition, and the mean ΔF/F0 ± SD was calculated from three replicates as described in the STAR Methods (left panel). SEpHluorin (green) and mCherry (red) signals were recorded, and the green-to-red ratio was converted to pH_i_ using a calibration curve (Figure S5A). The mean pH_i_ was calculated from three fields per condition, and these values were used to derive the mean pH_i_ ± SD based on three replicates (right panel). Statistical significance was assessed using two-way ANOVA with Tukey’s test (**p ≤ 0.01, ***p ≤ 0.001). (B) pH_lys_ was measured in pH_lys_ sensor cells pretreated with ML-SI3 (25 μM) for 30 minutes, followed by a 2-hour incubation in fresh media at pH 8.5 containing DMSO (control), or ML-SI3 (left panel). pH_lys_ was also measured over 2 h in pH_lys_ sensor cells transfected with either control siRNA or TRPML3- specific siRNA for 72 h before switching to fresh media at pH 8.5 (right panel). Quantification was based on mTFP1 and mCherry signals from 18–22 cells (left panel) or 27–35 cells (right panel) at each time point, across three independent replicates. Statistical significance was assessed using an unpaired two- tailed Student’s t-test (***p < 0.001). (C) Ca^2+^ levels were measured in Ca^2+^ sensor cells treated with either control siRNA or TRPML3 siRNA, followed by a 2-h incubation in media at pH 7.5 or pH 8.5. ΔF/F0 was measured in three fields per condition, and the mean ΔF/F0 ± SD was calculated from three replicates as described in the STAR Methods. Statistical significance was assessed using two-way ANOVA with Tukey’s test (***p < 0.001). (D, E) pH_i_ (D) and Ca^2+^ levels (E) were measured in sensor cells transfected with indicated siRNAs for 72 h, followed by a 2-h incubation with DMSO (control), ML-SA1 (25 μM) , or EBSS. Signal intensities were analyzed in three fields per condition, and the mean values were calculated from three replicates as described in the STAR Methods. Statistical significance was evaluated by two-way ANOVA with Tukey’s test (ns, not significant; *p < 0.05; ***p < 0.001). (F) Ca^2+^ levels in Ca^2+^ sensor cells (left panel) and pH_i_ in pH_i_ sensor cells (right panel) were measured after treatment with DMSO (control), SN-2 (30 μM), or ML-SA1 (25 μM) for 2 h. Signals were analyzed as described in panel A. Statistical significance was assessed using one-way ANOVA with Dunnett’s test (***p < 0.001). (G) Western blot of endogenous ARL8B and immunofluorescence analyses of LAMP1 in HeLa cells treated with DMSO (control), or SN2 (30 μM) for 2 h (left panel). Endogenous ARL8B protein levels were normalized, and relative levels are presented as mean ± SD (n = 3) (middle panel). Statistical significance was assessed using an unpaired two-tailed Student’s t-test (***p < 0.001). A SuperPlot depicting the ratio of perinuclear LAMP1 to total LAMP1, quantified using shell analysis of the immunofluorescence signals as described in Figure 1E (right panel). Horizontal lines represent mean ± SD (n = 3). Statistical significance was assessed using an unpaired two-tailed Student’s t-test (***p < 0.001). (H) Western blot analysis and quantification of endogenous ARL8B in HAP1 (WT, TRPML1 KO, and TRPML3 KO) cells treated with the indicated conditions for 2 h. Endogenous ARL8B protein levels were normalized, and relative levels are presented as mean ± SD (n = 3). (I) A model in which starvation, alkaline pH_e_, or ML-SA1 induces perinuclear lysosome accumulation by altering pH_i_ and Ca^2+^ levels, which in turn enhances RNF13-mediated degradation of ARL8B.

While low pH_lys_ enhances TRPML1 activity^62,63^, TRPML3 activity is stimulated by high pH_lys_^64,65^. To investigate the role of TRPML3 in the alkaline pH_e_-induced increase in cytosolic Ca^2+^, TRPML3 was silenced using siRNA, and subsequent changes in pH_lys_ were measured. In control siRNA-treated cells, alkaline pH_e_ caused an initial rise in pH_lys_ during the first 30 minutes, followed by a progressive decrease below the starting pH_lys_ levels over the course of a 2-h incubation (Figure 7B). Remarkably, in cells treated with ML-SI3 or depleted of TRPML3, the initial 30-minute increase in pH_lys_ was similar; however, pH_lys_ remained elevated for the subsequent 1.5 h. Furthermore, TRPML3 depletion inhibited the alkaline pH_e_-induced rise in cytosolic Ca^2+^ levels (Figure 7C). These findings suggest that alkaline pH_e_ likely activates TRPML3 by increasing pH_lys_. The subsequent Ca^2+^ release through TRPML3, activated by high pH_lys_, is presumed to enhance V-ATPase activity via ALG-2 activation, leading to a decrease in pH_lys_ (Figure S6D). While ML-SI3 effectively blocked the starvation-induced increase in pH_i_ (Figure 5E), it had only a minor effect on the pH_i_ increase caused by alkaline pH_e_ (Figure 7A). Since pH_i_ is influenced by both pH_e_ and V-ATPase activity and since the relationship between pH_i_ on pH_lys_, remains unclear, further investigation is needed to determine whether alkaline pH_e_ directly increases pH_lys_ through endocytosis of alkaline extracellular fluid or indirectly via changes in the pH_i_.

We next examined changes in pH_lys_ in TRPML1 or TRPML3 knockout (KO) cells treated with serum-free media, ML-SA1, or Torin1. In human HAP-1 wild-type cells, all three treatments lowered pH_lys_ (Figure S6E, F). Remarkably, the decrease in pH_lys_ by these stimuli was significantly impaired in TRPML3-KO cells, though less so than in TRPML1-KO cells. TRPML1 knockout or knockdown reduced the expression of both TRPML1 and TRPML3 proteins^66^, whereas TRPML3 knockout specifically affected TRPML3 expression (Figure S6G). RNF13 and ALG-2 levels were comparable across HAP-1 wild type, TRPML1-KO, and TRPML3-KO cells, whereas ARL8B levels were elevated in both TRPML1-KO and TRPML3-KO cells compared to wild-type cells (Figure S6H). These findings demonstrate that TRPML3 is crucial for proper lysosomal acidification in response to serum starvation, or treatment with Torin1 or ML-SA1. Since TRPML activation resulted in a concurrent decrease in pH_lys_ and an increase in pH_i,_ we sought to better understand the role of TRPML channels in regulating RNF13 activity. TRPML isoforms were depleted using siRNA and assessed their effect on pHi and cytosolic Ca^2+^ levels following EBSS or ML-SA1 treatments. Knockdown of either TRPML1 or TRPML3 inhibited the EBSS- or ML-SA1-induced increase in pH_i_ and cytosolic Ca^2+^ levels, with TRPML1 knockdown exerting a stronger inhibitory effect (Figure 7D, E).

Since the synthetic agonist ML-SA1 and antagonist ML-SI3 affect both TRPML1 and TRPML3 isoforms, the contribution of TRPML3 was further evaluated using the TRPML3-specific agonist SN-2. Treatment with SN-2 increased cytosolic Ca^2+^ levels and pH_i_ (Figure 7F), decreased ARL8B levels, and induced perinuclear lysosomal positioning (Figure 7G). These findings suggest a regulatory role for TRPML3 in lysosomal positioning. To further explore the relationship between TRPML channels and ARL8B protein levels, we examined ARL8B levels in TRPML1- or TRPML3-KO cells under conditions that activate TRPML channels. In HAP-1 wild-type cells, starvation, alkaline pH_e_, or treatment with ML-SA1 or SN-2 reduced ARL8B levels. However, these treatments failed to significantly decrease ARL8B levels in TRPML1-KO and TRPML3-KO cells (Figure 7H). Taken together, these findings suggest that TRPML3 plays a major role in regulating RNF13 activity and ARL8B levels in response to changes in nutrient availability and pH_e_. Based on all the findings, we propose a model in which starvation, alkaline pH_e_, or ML-SA1 induces perinuclear lysosome accumulation by altering pH_i_ and Ca^2+^ levels, which in turn enhances RNF13-mediated degradation of ARL8B (Figure 7I).

### The loss of ubiquitin ligase activity in RNF13 L312P is implicated in the pathogenesis of DEE-73

Missense variants in the *RNF13* gene, specifically the substitution of leucine at positions 311 or 312 with serine (L311S) or proline (L312P), have been implicated in the neurological disorder DEE-73 in humans^50^. To investigate the molecular mechanisms underlying DEE-73, we analyzed the expression levels, subcellular localization, and ubiquitin ligase activity of the RNF13 L311S and L312P mutants. Our findings revealed that the RNF13 L312P variant exhibited significantly higher expression levels compared to wild-type RNF13, whereas the RNF13 L311S variant showed expression levels similar to the wild type (Figure S7A-C). Autoubiquitination activity of RNF13 L312P was markedly reduced relative to both the wild type and RNF13 L311S (Figure S7B). Moreover, RNF13 L312P, but not RNF13 L311S, lost its ability to ubiquitinate and degrade ARL8B (Figure S7C-E).

We examined the function of the 307-EHTPLL-312 sequence, which resembles a [D/E]xxxL[L/I]-type lysosomal targeting signal, by analyzing the RNF13 ELL-AAA mutant with E307A, L311A, and L312A substitutions. The expression levels and ARL8B degradation activity of the RNF13 ELL-AAA mutant were comparable to those of the wild type (Figure S7C, E). Both the RNF13 L311S and RNF13 ELL- AAA mutants were primarily localized to lysosomes and induced perinuclear lysosomal localization similar to the wild type. In contrast, the ligase-dead RNF13 CS and L312P mutants failed to promote perinuclear lysosomal clustering and exhibited reduced co-localization with LAMP1. This reduction is likely attributable to impaired autoubiquitination (Figure S7F).

To further investigate the determinants of RNF13 localization, we generated chimeric RNF13 proteins fused with RNF128, another member of the PA-TM-RING E3 ligase family that predominantly localizes to the ER (Figure S7G). A chimera consisting of the N-terminal luminal and transmembrane domains of RNF128 with the C-terminal cytosolic domain of RNF13 was found to localize to the ER. Conversely, a chimera composed of the N-terminal RNF13 fused to the C-terminal RNF128 exhibited a localization pattern similar to RNF13, with substantial co-localization with LAMP1 (Figure S7G). This is consistent with previous findings that the PA domain is a key determinant for the endosomal localization of PA-TM-RING E3 ligases^67^. These findings suggest that the 307-EHTPLL-312 sequence of RNF13 does not function as a lysosomal targeting signal. Instead, the primary lysosomal localization signal resides within the N-terminal region of RNF13. Furthermore, the loss of ubiquitin ligase activity in the RNF13 L312P mutant likely contributes to the pathogenesis of DEE-73 in individuals with this variant. In contrast, the RNF13 L311S mutation does not impair ligase activity, suggesting that the associated defects may arise through other mechanisms. It is also possible that additional genetic factors contribute to the development of DEE-73.

## Discussion

ARL8B plays a critical role in regulating the intracellular positioning and functionality of lysosomes. In this study, we elucidate the mechanistic basis of ARL8B degradation triggered by alkaline pH_e_ or nutrient starvation. Our findings support a model in which pH_i_ and Ca^2+^, two key regulators of numerous cellular processes, modulate RNF13 activity to control lysosomal positioning in response to changes in pH_e_ or nutrient availability (Figure 7J).

### Regulation of lysosomal positioning by RNF13

RNF13 is a RING-type ubiquitin ligase that facilitates the transfer of ubiquitin directly from an E2 ubiquitin-conjugating enzyme to lysine residues on its substrates or itself. The protonation state of H332 in the PRR of RNF13 and the levels of Ca^2+^-activated ALG-2, which interacts with PRR, appear to regulate RNF13’s conformation. In our proposed model (Figure 6M), under acidic pH or low Ca^2+^ conditions, RNF13 adopts an inactive conformation. In this state, R217 in the NIR domain, critical for ARL8B binding, is predicted to form salt bridges with D291 and D293 in the NRR domain (Figure 2I), blocking ARL8B from accessing the substrate-binding site. Conversely, under alkaline pH and high Ca^2+^ levels, Ca^2+^-activated ALG-2 binds to the PRR, where H332 is deprotonated. ALG-2 binding induces an active conformation by disrupting the NIR-NRR interaction, freeing R217 to engage with ARL8B. Consequently, RNF13-mediated ubiquitination of ARL8B is enhanced, promoting lysosomal transport to the perinuclear region.

Additionally, we observed that the endogenous levels of RNF13 itself are influenced by H^+^ and Ca^2+^ levels. In its inactive conformation, restricted access of the E2-Ub complex to the RING domain or limited availability of target lysines on RNF13 may reduce its auto-ubiquitination. Interestingly, C-terminal deletion mutants of RNF13 that retain the NRR but lack the PRR exhibit severely reduced ARL8B degradation while displaying highly active autodegradation. These findings suggest that while H⁺ and Ca^2+^ levels modulate RNF13-mediated ubiquitination of both ARL8B and RNF13 itself, the interaction of ARL8B with the RNF13 substrate-binding site is the primary determinant of ARL8B ubiquitination. This underscores the critical role of RNF13 conformational changes, triggered by intracellular pH and Ca²⁺ fluctuations, in regulating ARL8B levels.

Recent studies have shown that SKIP induces removal of RAB7 from lysosomes by recruiting the cognate GTPase-activating protein TBC1D15^68^. Since SKIP is the primary effector of ARL8B, which drives the anterograde trafficking of lysosomes, an increase in ARL8B levels is expected to reduce the presence of RAB7-the key GTPase mediating retrograde trafficking through its interacting with RILP- on the same lysosomes. This shift in the ratio of ARL8B to RAB7 is presumed to amplify the effect of RNF13-mediated changes in ARL8B level on lysosomal trafficking. This mechanism may explain why small alterations in ARL8B levels result in disproportionately large effects on lysosomal positioning.

ALG-2 was shown to directly bind to TRPML1 in a Ca^2+^-dependent manner^69^ and to interact with dynamytin, a component of the dynein-dynactin complex independently of Ca^2+^, promoting retrograde trafficking of lysosomes^42^. Under acidic pH_i_ conditions, however, the activation of ALG-2 by ML-SA1 does not induce perinuclear positioning of lysosomes. Further studies are required to understand how TRPML1-ALG-2-dynein pathway is inhibited under acidic conditions.

During cancer development and progression, significant changes in the cellular distribution of lysosomes are observed^70^. The redistribution of lysosomes toward the cell periphery is critical for cancer cell growth, invasion, and metastasis^6^. This phenomenon has been documented in various cancers, including bladder^71^, breast^72^, and prostate cancers^73^. The acidic tumor microenvironment may play a role in driving this peripheral lysosomal distribution^74,75^. Notably, ARL8B has been identified as a key facilitator of lysosomal protease release and tumor invasion by promoting the peripheral positioning of lysosomes^72,73^. However, it remains unclear whether RNF13-dependent degradation of ARL8B is inhibited during cancer development. Further studies are necessary to explore this potential mechanism and its implications for cancer progression.

Changes in V-ATPase and TRPML channel activities can affect lysosomal concentrations of not only H^+^ and Ca^2+^ but also other ions due to their interconnectivity. Therefore, V-ATPase and TRPML channels are closely linked to lysosomal membrane potential and membrane tension. Although the influence of membrane potential on lysosomal trafficking is not well understood, it is known that high membrane tension inhibits membrane remodeling, while relief of tension promotes it^76^. Interestingly, a recent study suggested that TRPML2 activity is connected to membrane tension, while TRPML1 and TRPML3 are not responsive to osmolality changes^77^. Since we do not have any information on lysosomal membrane potential or membrane tension in our experimental conditions, we cannot exclude the possibility that these factors could influence lysosomal trafficking, potentially affecting the conformation and activity of lysosome membrane-bound RNF13.

### Regulation of pH_i_ by lysosomes

Cytosolic pH regulation is primarily governed by plasma membrane transporters^78^. However, the role of lysosomes in controlling pH_i_ remains largely unexplored. Despite occupying only 0.5–5% of the total cell volume, lysosomes maintain an H^+^ concentration 500- to 1,000-fold higher than that of the cytoplasm through the activity of the V-ATPase proton pump on their membranes^79^. Consequently, V-ATPase is predicted to regulate both pH_i_ and pH_lys_ simultaneously. Our study highlights the pivotal role of lysosomes in starvation-induced cytosolic alkalinization, which subsequently impacts their intracellular distribution. We further reveal that Ca^2+^ efflux through TRPML channels is tightly coupled to V-ATPase-driven H^+^ influx via activation of the Ca^2+^ sensor ALG-2. Previous studies showed that mTOR inhibition by Torin1 enhances V-ATPase assembly, reducing pH_lys_ and increasing lysosomal protease activity^60^. Consistent with this, we confirmed that Torin1 decreases pH_lys_ and found that it simultaneously raises pH_i_ and that these changes are dependent on TRPML channel activity and ALG-2. Under starvation conditions, V-ATPase assembly is known to increase^80^. Our findings suggest that starvation-induced lysosomal Ca^2+^ release and ALG-2 activation likely enhance V-ATPase assembly. Although Torin1 induces changes in pH_i_ and pH_lys_, similar to starvation or ML-SA1 treatment, it does not elevate cytosolic Ca^2+^ levels sufficiently to induce perinuclear lysosomal clustering. Interestingly, mTOR inhibition has previously been reported to activate TRPML1 and TRPML3^81–83^. How mTOR inhibition affects TRPML channels and ALG-2 remains to be fully elucidated and merit further investigation.

A recent study revealed that STAT3 associates with V-ATPase and enhances its activity. Notably, under acidic pH_i_ conditions, increased STAT3 translocation to lysosomes contributes to cytoplasmic neutralization by upregulating V-ATPase activity^84^. Our study further reinforces the crucial role of lysosomes in pH_i_ regulation. Together, these findings highlight emerging role of lysosomes as active participants in cellular pH homeostasis.

The reversible assembly of V_1_ and V_0_ subcomplexes is a key mechanism controlling V-ATPase activity. Several factors that regulate lysosomal distribution also influence V-ATPase assembly or activity. For instance, RILP interacts with the V1G1 subunit of V-ATPase, potentially modulating its activity^85^. TMEM55B has been shown to interact with V-ATPase, affecting its assembly^86^. Recent findings further suggest that nutrients dynamically regulate lysosomal function and positioning via the conversion of Phosphatidylinositol-3-phosphate (PI3P) to Phosphatidylinositol-4-phosphate (PI4P), which modulates V-ATPase assembly and activity^14^. PI4P acts as a pH sensor within the physiological pH range^87^, and its activation of V-ATPase likely changes PI4P’s protonation state, potentially affecting the binding affinity of effector proteins. These changes could further influence lysosome dynamics. Future research should explore whether these mechanisms of retrograde lysosomal trafficking during starvation also involve RNF13-mediated reduction of ARL8B, linking pH regulation to lysosomal repositioning.

### Role of TRPML3 in regulation of lysosomal positioning

Our study identifies TRPML3 as a key regulator of lysosomal Ca^2+^ release during starvation. While TRPML1 knockdown resulted in a more pronounced reduction in starvation-induced intracellular Ca^2+^ levels compared to TRPML3 knockdown, the precise contribution of TRPML1 to lysosomal Ca^2+^ release in this context remains unclear. This ambiguity arises from the simultaneous reduction in TRPML3 levels observed upon TRPML1 depletion. PI3P has emerged as an endogenous activator of TRPML3 during starvation^66^. Specifically, PI3P activates TRPML3 on the phagophore, facilitating autophagy induction. However, whether PI3P and/or phosphatidylinositol 3,5-bisphosphate (PI(3,5)P_2_), an endogenous agonist of both TRPML1 and TRPML3^13^, activates TRPML3 on lysosomes under starvation conditions remains to be clarified.

Our findings also underscore the critical role of TRPML3 in lysosomal pH homeostasis and positioning in response to alkaline pH_e_. TRPML3 is activated by elevated pH_lys_^64,65^. Alkaline pH_e_ raises lysosomal lumen pH, triggering TRPML3-dependent lysosomal Ca^2+^ release. This Ca^2+^ release activates ALG-2, which is thought to enhance V-ATPase assembly. Supporting this hypothesis, V-ATPase assembly was shown to increase upon treatment with pH_lys_-elevating lysosomotropic agents, including NH₄Cl, chloroquine, and L-leucyl L-leucine-O-methyl-ester (LLOMe)^80,88^. Additionally, our results suggest that TRPML3 functions as a sensor for pH_lys_, playing a pivotal role in maintaining lysosomal acidity. By activating V-ATPase in response to elevated pH_lys_, TRPML3 helps restore the acidic environment essential for optimal lysosomal function. This mechanism highlights the importance of TRPML3 in preserving lysosomal homeostasis and ensuring the proper functioning of these organelles.

### Role of ALG-2 in regulation of lysosomal positioning

In addition to activating RNF13, ALG-2 plays a critical role in starvation-induced perinuclear lysosomal clustering. This process involves the activation of V-ATPase, binding to TRPML1, and the recruitment of the dynein-dynactin motor complex. Although thapsigargin treatment induces elevated cytosolic Ca^2+^ levels, the absence of significant changes in pH_i_ suggests that increased Ca^2+^ alone is insufficient for ALG-2-mediated V-ATPase activation. Previous studies have indicated that ALG-2 recruitment to lysosomes requires its interaction with TRPML1 and localized elevations in perilysosomal Ca^2+42^. While TRPML1 has been shown to mediate local Ca^2+^ signals to regulate autophagy^47^ and lysosome positioning^89^, our study indicates that activation of TRPML1 leads to a global increase in Ca²⁺ levels, consistent with recent reports^90–92^. The observation that thapsigargin, in combination with Torin1, activates RNF13 suggests that under conditions of mTOR inhibition, ALG-2 may be further activated by a global rise in Ca^2+^ levels. However, the precise mechanisms by which ALG-2 responds to local versus global Ca²⁺ dynamics remain unclear. Further research is needed to delineate the molecular pathways through which ALG-2 modulates lysosomal function in response to Ca^2+^ signaling.

### Altered RNF13 activity in the pathogenesis of DEE-73

Our data demonstrate that the RNF13 L312P variant exhibits significantly impaired autoubiquitination and ARL8B ubiquitination activity. In contrast, the L312A substitution in the RNF13 ELL-AAA mutant retains ligase activity comparable to the wild-type protein. This suggests that the proline substitution at position 312 may induce a structural kink in the C-terminal region, potentially obstructing the access of ALG-2 to the PRR or E2 to the RING domain. Additionally, a recently reported nonsense variant (c.901G>T; p.Glu301X) associated with DEE-73^50^ likely retains the NRR but lacks the PRR, resembling RNF13 (1-305), which is incapable of ubiquitinating ARL8B. Since ARL8B is known to regulate neural connectivity during brain development by controlling axon branch positioning and density^26^, the inability of these RNF13 variants to regulate ARL8B levels may underlie, at least partially, the biological mechanisms contributing to DEE-73 symptoms.

In conclusion, this study sheds light on the regulation of retrograde and anterograde lysosomal transport in response to pH_e_ and nutrient availability. These findings advance our understanding of lysosomal heterogeneity, including luminal pH variations and subcellular localization. Although the SKIP- mediated coupling of lysosomes to kinesin-1 through interaction with ARL8B is a central mechanism of ARL8B-dependent anterograde lysosomal trafficking, several ARL8B effectors have also been identified^28,31–33^. However, their individual or cooperative roles under specific conditions remain poorly understood. Further research is warranted to explore the impact of RNF13 on ARL8B effectors and their involvement in lysosomal positioning and autophagy.

## Limitations of the study

This study has some limitations. Notably, we did not account for the heterogeneity of lysosomes or cytoplasmic regions. A more detailed analysis of pH and Ca^2+^ levels in individual lysosomes and specific cytoplasmic domains could provide deeper insights into the regulation of lysosome positioning. The pH_i_ sensor used in this study measures the average cytosolic pH, which may not accurately reflect local pH changes near lysosomes. Localized pH near lysosomes is likely influenced by V-ATPase activity following lysosomal Ca^2+^ release, potentially impacting signaling molecules situated on the lysosomal membrane. For instance, mTORC1 activity is reduced on perinuclear lysosomes compared to peripheral lysosomes^41^, possibly due to differences in local pH depending on their intracellular location.

## Acknowledgements

This work was supported by grants from the National Research Foundation of Korea (NRF- 2021R1A2C1011189) and Ministry of Science and ICT (RS-2023-00273665) by the Korean government. We thank to Young Jin Yoon from the Research & Technology Support Team, Songeui Medical Campus, The Catholic University of Korea, for technical assistance.

## Author contributions

1. L. T. T., A. V. V. , and Y. H. formal analysis; L. T. T., A. V. V., S. S., C. L., S-H. P., and E.-B. C. investigation; H. J. K. resources; L. T. T., and J.-B. Y. methodology; S. K. and J.-B. Y. conceptualization; S. K. and J.-B. Y. supervision; S. K., J.-B. Y., and H. J. K. writing-review and editing; S. K. and J.-B. Y. funding acquisition.

## Declaration of interests

The authors declare no conflict of interest.

## STAR methods

Key resources table

| REAGENT or RESOURCE | SOURCE | IDENTIFIER |
| --- | --- | --- |
| **Antibodies** |  |  |
| Anti-ARL8B | ThermoFisher Scientific | PA5-100124 |
| Anti-RNF13 | Cusabio | CSB-PA019831DSR1HU |
| Anti-FLAG | Sigma | F7425 |
| Anti-HA | Santa Cruz Biotechnology | sc-7392 |
| Anti-β-actin | GeneTex | GT5521 |
| Anti-ALG-2 | Santa Cruz Biotechnology | sc-376950 |
| Anti-GADPH | Santa Cruz Biotechnology | sc-166574 |
| Anti-LC3 | MBL | PM036 |
| Anti-p62 | Novus Biologicals | H00008878 |
| Anti-Ub | Santa Cruz Biotechnology | sc-166553 |
| Anti-ARL8B | Proteintech | 13049-1-AP |
| Anti-TRPML1 | Alomone Labs | ACC-081 |
| Anti-TRPML3 | Alomone Labs | ACC-083 |
| Anti-Lamp1 | Santa Cruz Biotechnology | sc-18821 |
| **Chemicals, peptides, and recombinant proteins** |  |  |
| Lipofectamin 2000 | Invitrogen | 11668500 |
| Lipofectamine RNAiMax | Invitrogen | 13778150 |
| MES | Sigma | M8250 |
| HEPES | Sigma | H6147 |
| ML-SA1 | Sigma | SML0627 |
| Torin 1 | Cayman Chemical | CAY10997 |
| ML-SI3 | MedChemExpress | HY-139426 |
| Penicillin-streptomycin | GenDEPOT | CA005-010 |
| Blasticidin | InvivoGen | ant-bl-05 |
| Puromycin | Enzo Life Sciences | BML-GR312-0050 |
| Hygromycin B | InvivoGen | ant-hg-1 |
| SN-2 | MedChemExpress | HY-16696 |
| Nigericin solution | Sigma | SML1997 |
| Thapsigargin | Sigma | T9033 |
| **Oligonucleotides and Recombinant DNA** |  |  |
| Primers used for cloning and site-directed mutagenesis, and oligonucleotides used for complex reconstitution are listed in Table S1 | This paper | Table S1 |
| **Software and algorithms** |  |  |
| Prism 8 | GraphPad | https://www.graphpad.com/scientific-software/prism/ |
| Fiji/ImageJ | Fiji/ImageJ | N/A |
| GalaxyWEB | Ko et al.<sup>93</sup> | https://galaxy.seoklab.org/ |
| COOT | Emsley et al.<sup>94</sup> | https://www2.mrc-lmb.cam.ac.uk/personal/pemsley/coot/ |
| Biorender | Biorender | https://www.biorender.com/ |
| Microsoft Excel | Microsoft | https://www.microsoft.com/ |
| Others |  |  |
| DMEM | Welgen | LM001-50 |
| EBSS | Welgen | LB002-03 |
| Lysosensor DND-189 | Invitrogen | L7535 |
| FBS | GenDEPOT | F0900-050 |
| anti-FLAG M2 affinity gel | Sigma | A2220 |
| Dynabeads® Protein G | Thermo Fisher Scientific | 10003D |
| Ubiquitin IP Assay | Cytoskeleton, Inc | BK161 |
| DPBS | Welgen | LB 001-20 |
| Iscove's modified Dulbecco's medium | Sigma | I3390-500ML |
| SPL confocal dish | SPL Life Sciences, | 100350 |
| RNAiso Plus | Takara | 9109 |
| cDNA Synthesis Kit | Takara Bio | 6210 |
| TB Green Premix Ex Taq | Takara Bio | RR420A |

## Resource availability

### Lead contact

Further information and requests for resources and reagents should be directed to and will be fulfilled by the lead contact, Jong-Bok Yoon.

## Material availability

Plasmids and cell lines generated in this study are available from the lead contact upon request.

## Data and code availability

- Data reported in this paper will be shared by the lead contact upon request.
- This paper does not report original code.
- Any additional information required to reanalyze the data reported in this paper is available from the <u>lead contact</u> upon request.

Experimental model and study participant details

## Cell culture

HeLa, HEK293T, and MCF7 cells were cultured in high-glucose Dulbecco’s modified Eagle’s medium (DMEM) (WelGENE) supplemented with 10% fetal bovine serum (FBS) (GenDEPOT) and 1% penicillin/streptomycin at 37 °C with 5 % CO_2_. HAP-1 (TRPML1-KO, TRPML3-KO, and WT) cells were purchased from Horizon and cultured in Iscove’s modified Dulbecco’s medium (Sigma) supplemented with 10% FBS, 4 mM L-Glutamine, 1 mM sodium pyruvate, and 1% antibiotics. A cell line expressing mRFP-GFP-LC3, was a kind gift from Dr. M. Komatsu (Juntendo Univ) ^95^, and cells were cultured in DMEM supplemented with 10 % FBS and 1 % penicillin/streptomycin.

## Method details

### Plasmids

The clones for the complementary DNAs (cDNAs) encoding human *ARL8B* (accession no. NM_018184.2) and *RNF13* (accession no. NM _007282.4) were obtained from DNASU Plasmid Repository. The plasmid for the expression of epitope-tagged *ARL8B* was constructed by subcloning the PCR-amplified cDNA into the in-house modified pcDNA3.1 (Invitrogen), which has a C-terminal FLAG- or HA-tag as described^25^. The full-length complete coding sequence of *RNF13* and the deletion mutants for RNF13 were generated by subcloning each PCR-amplified cDNA into the modified pcDNA3.1 or pYR, generating plasmid constructs with the C-terminal FLAG, or HA-tag using ClaI and XbaI restriction sites. To generate RNF13 Δ205-237, two specific fragments of RNF13 including 1-204, and 238-381 were generated with unique primers: fragment 1-204 with RNF13 FL sense primer and RNF13 1-204 antisense-1 primer, and fragment 238-381 with RNF13 238-381 sense primer and RNF13 FL antisense primer. To generate RNF13 Δ205-219, two specific fragments of RNF13 including 1-204 and 220-381 were generated: fragment 1-204 with RNF13 FL sense primer and RNF13 1-204 antisense- 2 primer, and fragment 220-381 with RNF13 220-381 sense primer and RNF13 FL antisense primer. To generate RNF13 Δ220-233, two specific fragments of RNF13 including 1-219 and 234-381 were generated with unique primers: fragment 1-219 with RNF13 FL sense primer and RNF13 1-219 antisense primer, and fragment 234-381 with RNF13 234-381 sense primer and RNF13 FL antisense primer. After generating the two specific fragments, they were joined using overlap extension PCR with the RNF13 FL primer set. To generate RNF13N/RNF128C, the N-terminal of RNF13 was fused to the C-terminal of RNF128, and RNF128N/RNF13C were generated by fusing N-terminal of RNF128 with the C-terminal of RNF13. The point mutation constructs including C243S, R210,214A, R215,217A, K220A, K220,224A, K220,224,233A (3KA), D291,293,295,297A (4DA), H308A, H332A, H332K, H361A, L311S, L312P, L311,312A, and E309,L311,L312A were generated using QuikChange Kit following the manufacturer’s protocol (Stratagene). To generate the plasmid expressing siRNA-resistant full-length RNF13, we introduced three single-nucleotide mutations (T300G, T303C, and T306C) in the coding sequence without causing any changes in amino acids (sense mutation) using QuikChange Kit (Stratagene). The ARL8B mutant constructs, K131R, K141R, and K146R, were generated using the QuikChange Kit according to the manufacturer’s protocol (Stratagene). The cDNA clone for ALG-2 was obtained from the Korean Human Genebank (hMU001447). The epitope-tagged ALG-2 was generated by subcloning the PCR-amplified cDNA into the modified pcDNA3.1 with C-terminal FLAG or HA tags using BgIII and XhoI sites. The ALG-2 mutant constructs, E47A or E114A, were generated using QuikChange Kit following the manufacturer’s protocol (Stratagene). To generate the ARL8B-mCherry construct, the fusion construct of the ARL8B and mCherry was first created by the ARL8B cDNA and the mCherry cDNA from the pFUGW-FIRE-pHLy plasmid (Addgene #170774)^59^ via jumping PCR amplification. Then, the resulting PCR product was subcloned into the pCDH-CMV-MCS-EF1-Hygro vector using NheI and BamHI restriction enzymes. To generate the LAMP1-GFP fusion construct, the LAMP1 cDNA was first inserted into the pEGFP-N1 vector using NheI and AgeI restriction sites. The LAMP1-GFP fusion construct was then cloned into the pCDH-CMV-MCS-EF1-Neo vector using NheI and BamHI restriction sites. All constructs were confirmed by sequencing. The primer sequences used for cloning and the vectors cloned into are listed in Table S1.

## RNAi

SiRNAs against RNF13, ARL8B, ALG-2, RNF167, TRPML1, and TRPML3 were generated (Bioneer) and transfected using Lipofectamine RNAiMax reagent (Invitrogen) according to the manufacturer’s instructions. For knockdown experiments, cells were transfected with 60 nM siRNA for 72 h for RNF13, ARL8B, ALG-2, and RNF167. For knockdown of TRPML1 and TRPML3 expression, cells were transfected with 60 nM siRNA for 24 h, followed by 20 nM siRNA transfection for another 48 h. The sense siRNA sequence were as follows: siRNF13#1, 5’-UUAGAAGACUUGAUUGUAA-3’; siRNF13#2, 5’-GAAACUUCCUGUACAUAAA-3’; siRNF13#3, 5’- GCCACCUUAUCUUAGUUCCAG-3’; siARL8B, 5’-AGGUAACGUCACAAUAAAGAU-3’; siALG- 2, 5’-AAAGACAGGAGUGGAGUGAUAUCAG-3’; siRNF167, 5′- UAGCUCGUUGUAUCCAGCACCGGAA-3′; siTRPML1, 5’-CCCACAUCCAGGAGUGUAA-3’; and siTRPML3, 5’-GGAUGGUACAUUAUGAUUAUU-3’.

## RNA extraction and Quantitative real-time PCR (qRT-PCR)

Total RNA was extracted from cells using Trizol Reagent (Invitrogen, Carlsbad, USA) according to the manufacturer’s protocol. Complementary DNA (cDNA) was synthesized using a cDNA synthesis kit (Takara, Shiga, Japan). qRT-PCR was performed using an SYBR Green qRT-PCR analysis kit (Takara, Shiga, Japan). Messenger RNA (mRNA) levels of RNF167 were quantified using the 2-^ΔΔ^Ct method, with GAPDH mRNA as the reference gene. Primers for RNF167 are as follows: 5’-TCC AGG GGT TCC TTG TGG A-3’ (sense) and 5’-CCT TCT GGG CAT TTA GGA CCT T-3’ (antisense). Primers for GAPDH are as follows: 5’-AACTTTGGCATTGTGGAAGG-3’ (sense), and 5’- ACACATTGGGGGTAGGAACA-3’ (antisense).

## Cell line generation

To generate RNF13 or RNF167 knockdown cell lines, HEK293T cells were co-transfected with psPAX2, PMD2.G and either pLKO.1-shRNF13-puro or pLKO-shRNF167-puro using Lipofectamine 2000 transfection reagent. The pLKO.1-shRNF13-puro and pLKO.1-shRNF167-puro plasmids were constructed by subcloning the oligonucleotides listed in Table S1 into pLKO.1 puro (Addgene, #10878). Lentivirus-containing media were collected 72 h post-transfection and used for the transduction of HeLa and MCF7 cells in the presence of 10 μg/mL polybrene (Sigma). The medium was changed to complete DMEM supplemented with 1 µg/mL puromycin at 72 h post-transduction. Selection continued for 2 days until complete infection was achieved. The cells were cultured and used for subsequent experiments.

To generate HeLa cells stably expressing both ARL8B-mCherry and LAMP1-GFP, cells were first transduced with lentivirus containing the plasmid encoding *ARL8B-mCherry* for 72 h. After selection with hygromycin continued for additional 5 days, the cells were transduced with lentivirus carrying the plasmid encoding *LAMP1-GFP*. Selection was continued with G418 for an additional 5 days until stable expression was achieved.

The sensor cell lines for relative measurement of pH_i_, pH_lys_, and cytoplasmic Ca^2+^ levels were generated in a HeLa cell background. For measuring pH_i_, pCDH-Hygro-mCherry-SEpHluorin plasmid was constructed by subcloning the DNA fragment encoding mCherry-SEpHluorin (Addgene, #32001)^57^ into pCDH-CMV-MCS-EF1 α-Hygro (System Biosciences, #CD515-1) using XbaI and BamHI sites. This construct includes mCherry and SuperEcliptic (SE) pHluorin, which are pH-insensitive and pH-sensitive, respectively. Cell lines stably expressing mCherry-SEpHluorin (pH_i_ cell line), FIRE-pHLy (pH_lys_ cell line), or GCaMP6f (cytosolic Ca^2+^ cell line) were generated by lentiviral transduction. HEK293T cells were transfected with pCDH-Hygro-mCherry-SEpHluorin, pFUGW-FIRE-pHLy (Addgene, #170774)^59^, or pLX304-GCaMP6f (Addgene, #163045)^61^ with psPAX2, and PMD2.G using Lipofectamin 2000 for 72 h. The lentivirus was transduced into HeLa cells in the presence of 10 μg/mL polybrene for 72 h. Cells were selected using 200 µg/mL hygromycin B for the pH_i_ cell line and 10 µg/mL blasticidin for the cytosolic Ca^2+^ cell line. For the pH_lys_ cell line, cells were used for the subsequent experiments immediately after a 72-h transduction with lentiviral medium without selection, due to the high toxicity of zeocin to the transduced cells.

## Cell transfection and treatment

Lipofectamine RNAiMax (Invitrogen) or polyethylenimine (Sigma) were used for the transfection of cell lines. MG132 and Torin1 were purchased from Caymen Chemical Company, Bafilomycin A1, ML-SA1, ML-SI3, and thapsigargin from Sigma, and SN-2 from MedChemExpress.

For starvation experiments, cells were cultured in complete media for 48 h before undergoing starvation, following previously described protocols^41^. For starvation experiments, two different conditions were used. For complete nutrient deprivation, cells were washed with phosphate-buffered saline (PBS) and incubated in EBSS (Welgen) at 37°C with 5% CO_2_ for the duration specified in the figure legends. For milder starvation, cells were washed with PBS and incubated in serum-free DMEM at 37°C with 5% CO_2_ for the indicated duration in figure legends.

To establish stable acidic or alkaline pH conditions, media formulations were prepared as described in a previous study^96^. DMEM (Gibco) containing 25 mM glucose and 4 mM L-glutamine was supplied with 5 % FBS, 0.25× penicillin-streptomycin, and 25 mM MES (pKa 6.66 at 37°C, useful pH range of 6.1– 7.5 at 25 °C, Sigma), or 25 mM HEPES (pKa 7.31 at 37 °C, useful pH range of 6.8–8.2 at 25 °C, Sigma), or 25 mM Tris (pKa 7.72 at 37 °C, useful pH range of 7.2–9.0 at 25 °C, Sigma). Media were adjusted to pH 6.5, 7.5, or 8.5 using NaOH, or HCl before filter sterilization and used in atmospheric CO_2_.

## Identification of ARL8B as a potential RNF13 substrate using proximity-dependent biotin labeling

HeLa cells were transfected with the constructs encoding RNF13-BirA-FLAG and AP-HA-Ub for 24 h and then treated with 50 μM Biotin and 25 μM MG132 for 4 h. Purification of the biotinylated– ubiquitinated proteins was performed as described previously ^54,97^. Briefly, cells were harvested and lysed in the homogenization buffer (0.25 M sucrose, 10 mM HEPES, pH 7.4, 1 mM EDTA, 10 mM PMSF, and 10 mM NEM). Cell lysates were centrifuged for 60 min at 28000 x*g* to collect membrane-enriched fraction, which was subsequently solubilized in lysis buffer (2% SDS, 250 mM NaCl, 50 mM Tris-Cl, pH 7.4). The lysates were boiled at 95**°**C for 10 min, centrifuged for 10 min at 12000 x*g*, and incubated with anti-FLAG M2 affinity gel overnight at 4 **°**C. The lysates containing biotinylated proteins were incubated with streptavidin agarose in a buffer (250 mM NaCl, 50 mM Tris-Cl, pH 7.4) overnight at 4 **°**C. Subsequently, beads were reduced, alkylated, and trypsinized into peptides. Finally, trypsinized peptides were purified using ubiquitin branch motif (K-ε-GG) immunoaffinity beads (Cell Signaling Technology) and analyzed using nanoelectrospray LC-ML/MS on an LTQ Orbitrap Velos (Thermo Fisher Scientific) as described previously ^54,97^. For matching proteins using MS/MS spectra in Mascot (Matrix Science), the following parameters were used: 2 missed cleavage sites was chosen, carbamidomethyl (C) as fixed modification, monoisotopic for mass value, protein mass unrestricted, 10 ppm for peptide mass tolerance, and 0.8 Da for fragment mass tolerance.

## Co-immunoprecipitation and Western blot analysis

HeLa cells were cultured for 24 h post-transfection with the indicated plasmids, then harvested by trypsinization, followed by washing and centrifugation. Cells were immediately placed in lysis buffer [50 mM Tris-HCl (pH 7.4), 150 mM NaCl, 1 mM EDTA, 1% Nonidet P-40, 1 mM dithiothreitol, and 0.2 mM PMSF] for 30 min on ice. Cell lysates were either analyzed directly by western blot analysis or processed further for co-immunoprecipitation (Co-IP) experiments. Co-IP was conducted as follows: lysates were centrifuged at 21000 x*g* for 15 min at 4 °C; the supernatants were then incubated with anti-FLAG M2 affinity gel (Sigma) overnight at 4 °C; after brief centrifugation at 1500 rpm, the pellets were washed with lysis buffer containing 0.5% NP40 and the immunoprecipitated proteins were eluted by incubating the pellets with SDS sample buffer (200 mM Tris-Cl (pH 6.8), 2% SDS, 0.4% bromophenol blue, and 40% glycerol) for 30 min at 37 °C. For immunoprecipitation with anti-ALG-2 or anti-RNF13 antibodies, cell lysates were incubated with the antibodies in the presence of 5 mM EGTA or 100 μM CaCl_2_ overnight at 4 °C. The lysates were then incubated with Dynabeads® Protein G for 2 h at 4 °C. The beads were washed with lysis buffer and subsequently processed for western blot analysis. Protein samples were separated by SDS-polyacrylamide gel electrophoresis, transferred onto nitrocellulose membranes (Pall Corp., Pensacola, FL), and subjected to western blot analysis. The band was visualized using the ECL system (DonginBiotech, Seoul, Korea).

## Ubiquitination assay

HeLa cells were transfected with a plasmid encoding T7-ubiquitin along with the expression constructs specified in the figure legends. After 24 h, the transfected cells were treated with 25 μM MG132 for 6 h. The cells were then harvested and lysed using a lysis buffer. The lysates were centrifuged at 21000 x*g* for 15 min at 4 °C to remove cell debris. The resulting supernatants were incubated with anti-FLAG M2 affinity gel (Sigma) overnight at 4 °C. After centrifugation, the pellets were washed with lysis buffer, eluted in SDS sample buffer, and analyzed by western blot analysis.

Ubiquitination of endogenous ARL8B was analyzed using the SignalSeeker Ubiquitination Detection Kit (Cytoskeleton) following the manufacture’s instruction. Briefly, HeLa cells were transfected with the indicated constructs or treated with the specified conditions as described in the figure legends. Cell lysates were washed, incubated with BlastR lysis buffer (with NEM and protease inhibitor), passed through a BlastR filter, and diluted with BlastR dilution buffer. They were then incubated with control or ubiquitination affinity beads for 2 h at 4°C, washed with BlastR-2 buffer, and eluted with bead elution buffer. Finally, samples were collected using spin columns and processed for western blot analysis.

## Immunofluorescence microscopy

Cells grown on coverslips were washed with PBS and fixed in 4% paraformaldehyde for 15 min. Cells were permeabilized with 0.5% PBS-T (0.5% Triton X-100 in PBS) for 5 min, washed with 0.1% PBS-T, and blocked with 10% normal goat serum for 30 min. Samples were incubated with primary antibody overnight at 4 °C. Cells were washed three times with 0.1% PBS-T before incubating with fluorescently conjugated secondary antibody for 50 min. Finally, cells were washed with 0.1% PBS-T, and nuclei were stained with DAPI. Immunofluorescence images were obtained using a confocal laser scanning microscope (LSM 700 or LSM800; Zeiss, Gottingen, Germany) or an inverted fluorescence microscope (LSM510; Carl Zeiss, Germany).

## Time-lapse confocal imaging

HeLa cells stably expressing ARL8B-mCherry and LAMP1-GFP were seeded onto an SPL confocal dish. Time-lapse fluorescence imaging was performed using an Olympus FV3000 scanning confocal microscope equipped with a 60×1.42 NA oil immersion objective at room temperature. Cells were incubated in a HEPES-buffered media, washed twice with PBS, and placed under the specified conditions. A 488 nm Diode laser was used to excite GFP, and fluorescence emission was collected between 500– 540 nm using a GaAsP PMT detector in analog mode. Similarly, a 561 nm Diode laser was used to excite mCherry, with emission collected between 570–620 nm using the same detector. A series of sequential images at 12 bits were collected and images were obtained over 1 h period with a frame time of 1 minute (N = 60).

Fluorescence signal intensity was quantified by measuring mCherry signals in the cell undergoing time- lapse imaging every minute for 1 h using ImageJ. The relative signal for plotting was calculated as the decrease from the initial signal, normalized to the original value.

## Analysis of intracellular distribution of lysosomes

To analyze lysosome distribution, we used a “shell analysis” method^98^, utilizing confocal images acquired with a Zeiss LSM800 inverted laser scanning microscope (Carl Zeiss). The images were processed using FIJI as described previously ^31,98,99^. For accurate quantification, we selected cells with a relatively round and uniform shape, excluding narrow, elongated cells due to potential inaccuracies in analysis. These criteria were predefined and applied consistently across all conditions.

Cell boundaries were determined using the CD147 marker, which localizes both intracellularly and at the plasma membrane, while nuclear boundaries were defined by the DAPI signal. The LAMP1 signal from neighboring cells was excluded, and the total whole-cell LAMP1 signal was quantified. The nuclear outline was then expanded in fixed increments five times to create five concentric shells (areas). The LAMP1 signal was determined in each shell. Area 1 was defined as the perinuclear region of the cell, while Areas 4 and 5 were defined as the peripheral edge. Each cell was analyzed individually. For each condition, three independent experiments were conducted, with more than 20 cells per experiment. The ratio of perinuclear lysosomes was calculated as the LAMP1 signal in Area 1 divided by the total LAMP1 signal. Quantifications were confirmed by a researcher blinded to the experimental groups.

## Measurement of pH_i_ and pH_lys_

The pH_i_ was assessed using HeLa cells stably expressing mCherry-SEpHluorin (pH_i_ sensor cell line), which contains the pH-sensitive fluorescent protein SEpHluorin (green) fused with the pH-insensitive mCherry (red). A standard curve was generated by determining the green to red signal intensity ratio at various pH. Briefly, cells were washed twice with PBS and incubated in a high-potassium buffer to prevent a [K^+^] gradient from driving a proton gradient ^100,101^. The buffer contained (in mM): KCl 100, NaCl 38, CaCl_2_ 1.8, MgSO_4_ 0.8, and NaH_2_PO_4_ 0.9, with Good buffer (25 mM) pre-adjusted to pH 8.0, 7.5, 7.0, 6.5, and 6.0, along with 10 μm nigericin (Invitrogen). The Good buffer was chosen based on its pKa with MES for pH 6.5; HEPES for pH 7.0 and pH 7.5; and Tris for pH 8.0 and pH 8.5. Cells were cultured in complete DMEM for 48 h, then washed twice with PBS and treated as indicated in the figure legends. Images of 20× fields were captured using an Olympus IX70 inverted fluorescence microscope. Before calculating the SEpHluorin/mCherry ratio, images were corrected for background and shading using a BaSiC tool in the Fiji plugin ^102^. Three replicates per experiment, with at least three fields analyzed per replicate, were used to calculate the mean pH_i_ ± SD.

The pH_lys_ was assessed using HeLa cells stably expressing mTFP1–hLAMP1–mCherry. This construct contains a pH-sensitive mTFP1 fused to the lysosomal lumen-facing N-terminal portion of human LAMP1, and a pH-insensitive mCherry fused to the C-terminal of hLAMP1, facing outside the lysosome^59^. A standard curve was generated as described previously. The buffer composition included (in mM): NaCl 5, KCl 115, MgSO_4_.7H_2_O 1.3, and MES 25, pre-adjusted to a pH range of 3.0–6.0, along with 10 μM nigercin. Images of 20× fields were captured using an Olympus IX70 inverted fluorescence microscope. Cell boundaries were determined using the free hand selection tool in Fiji. The signals of mTFP1 (green) and mCherry (red) were measured by subtracting the background signal from the whole-cell signal. The ratio of mTFP1/mCherry was then calculated. At least 20 representative cells with three replicates were analyzed.

## Measurement of lysosomal pH using Lysosenser staining

For the lysosensor assay, cells were plated overnight in a 35-mm confocal cell culture dish or 96-well black plate. The following day, cells were treated with different conditions as described in the figure legends, and 2 μM Lysosensor DND-189 (Invitrogen) was applied for 1 h. To generate a pH standard calibration curve, cells were incubated for 10 min at 37 °C in sodium acetate-acetic acid calibration buffers (from pH 4 to pH 5.5). To clamp lysosomal pH, 10 μM nigericin and 1 mM KCl were added. The fluorescence of the lysosensor was observed with a confocal microscope (LSM800W/Airys-can, Carl Zeiss), and the fluorescence intensity was quantified with a fluorometer (Bio-Tek, Winooski, USA).

## Measurement of cytoplasmic Ca^2+^ levels

HeLa cells stably expressing GCaMP6f were grown for 48 h and incubated under different conditions as described in the figure legends. The images of 20× fields were captured using an Olympus IX70 inverted fluorescence microscope and corrected for background using Fiji. The fluorescence intensity of GCaMP6f (F) was monitored. Results are presented as ΔF/F0, where F0 is the baseline fluorescence and ΔF is the changes in fluorescence (ΔF = F − F0). Three replicates per experiment, with at least three fields analyzed per replicate, were used to calculate the mean ΔF/F0 ± SD.

## Molecular docking and refinement

The crystal structure of the RNF13 RING domain (PDB entry: 5ZC4) and the primary structure of the RNF13 acidic-rich region (aa. 291–305) were prepared for molecular docking. Molecular docking was conducted using GalaxyWEB (https://galaxy.seoklab.org/) ^93^. The RNF13 helical region (aa. 205–215) was extracted from the AlphaFold structure^56^. The connection between them was established using COOT^94^, followed by refinement using GalaxyWEB.

## Quantification and statistical analysis

GraphPad Prism v8.8 (San Diego, CA) and Microsoft Excel were used for calculations, analyses, and data visualization. Data are presented as mean ± SD. Statistical significance was assessed using an unpaired Student’s t-test, one-way analysis of variance (ANOVA), or two-way ANOVA, as detailed in the corresponding figure legends. Figures 1E, 1H, 4A, and others feature SuperPlots. These plots display individual data points as small circles, the means of each independent experiment as large, color-coded circles, and the mean ± SD of these means as a bar. Differences were considered statistically significant at *p < 0.05, **p < 0.01, or ***p < 0.001; "ns" denotes no significant difference. Quantifications and statistical analyses were based on three independent replicates, with over 20 cells observed in each replicate.

## Notes

### Competing Interest Statement

The authors have declared no competing interest.

