## Supplemental Figures for "RNF13 mediates pH- and Ca^2+^-dependent regulation of lysosomal positioning"

**Video S1. Time-lapse confocal imaging of a cell expressing ARL8B-mCherry and LAMP1-GFP placed in a medium at pH 7.5, related to Figure 1**

HeLa cells stably co-expressing ARL8B-mCherry and LAMP1-GFP were cultured in a medium at pH 7.4. For time-lapse imaging, cells were placed in a fresh medium at pH 7.5 and imaged 1-hour period using a 60× 1.42 oil immersion objective, as described in the STAR Methods. Images were acquired at 1-minute intervals.

**Video S2. Time-lapse confocal imaging of a cell expressing ARL8B-mCherry and LAMP1-GFP placed in a medium at pH 8.5, related to Figure 1**

HeLa cells stably co-expressing ARL8B-mCherry and LAMP1-GFP were cultured in a medium at pH 7.4. For imaging, cells were transferred to a fresh medium at pH 8.5 and analyzed as described in Video S1. Compared to the control (Video S1), cell exposed to an alkaline $pH_e$ exhibited a gradual perinuclear redistribution of lysosomes. Additionally, a reduction in ARL8B-mCherry fluorescence intensity was observed, which was further validated by immunoblotting and quantification (Figure S3G).

**Video S3. Time-lapse confocal imaging of a cell expressing ARL8B-mCherry and LAMP1-GFP placed in a serum-free medium, related to Figure 1**

HeLa cells stably co-expressing ARL8B-mCherry and LAMP1-GFP were cultured in a serum-containing medium, then transferred to a serum-free medium and imaged as described in Video S1. Compared to the control (Video S1), a cell in the serum-free medium exhibited a gradual perinuclear redistribution of lysosomes and reduced ARL8B-mCherry fluorescence, confirmed by immunoblotting and quantification (Figure S3G).

**Video S4. Time-lapse confocal imaging of a cell expressing ARL8B-mCherry and LAMP1-GFP treated with a negative control siRNA and placed in a medium at pH 8.5, related to Figure 1**

HeLa cells stably co-expressing ARL8B-mCherry and LAMP1-GFP were transfected with control siRNA for 72 h. For time-lapse imaging, cells were then transferred to a fresh medium adjusted to pH 8.5 and images were acquired every minute for 1 h as shown in Video S1. Cell transfected with control siRNA showed gradual perinuclear redistribution of lysosomes as seen in a cell treated with alkaline $pH_e$ (Video S2).

**Video S5, 6. Time-lapse confocal imaging of a cell expressing ARL8B-mCherry and LAMP1-GFP treated with an RNF13-specific siRNA and placed in a medium at pH 8.5, related to Figure 1**

HeLa cells stably co-expressing ARL8B-mCherry and LAMP1-GFP were transfected with an RNF13-specific siRNA for 72 h, then transferred to a fresh medium at pH 8.5 and imaged every minute for 1 h as shown in Video S1. Compared to the control siRNA transfected cell (Video S4), RNF13-depleted cells showed impaired alkaline media-induced lysosomal redistribution to the perinuclear region.

**Video S7. Time-lapse confocal imaging of a cell expressing ARL8B-mCherry and LAMP1-GFP treated with an RNF167-specific siRNA and placed in a medium at pH 8.5, related to Figure 1**

HeLa cells stably co-expressing ARL8B-mCherry and LAMP1-GFP were transfected with RNF167-specific siRNA for 72 h, then transferred to fresh medium at pH 8.5 and imaged every minute for 1 h as described in Video S1. Compared to the RNF13-depleted cells (Video S5, S6), RNF167-depleted cell did not show severe inhibition of alkaline $pH_e$-induced perinuclear lysosomal positioning nor reduction in ARL8B-mCherry fluorescence (Figure S3K).

**Video S8. Time-lapse confocal imaging of a cell expressing ARL8B-mCherry and LAMP1-GFP upon ML-SA1 treatment, related to Figure 4**

HeLa cells stably co-expressing ARL8B-mCherry and LAMP1-GFP were cultured in a complete medium, then transferred to a medium containing ML-SA1 (25 μM) and imaged every minute for 1 h as shown in Video S1. ML-SA1 treatment induced perinuclear lysosomal accumulation.

**Supplementary Figure 1.**

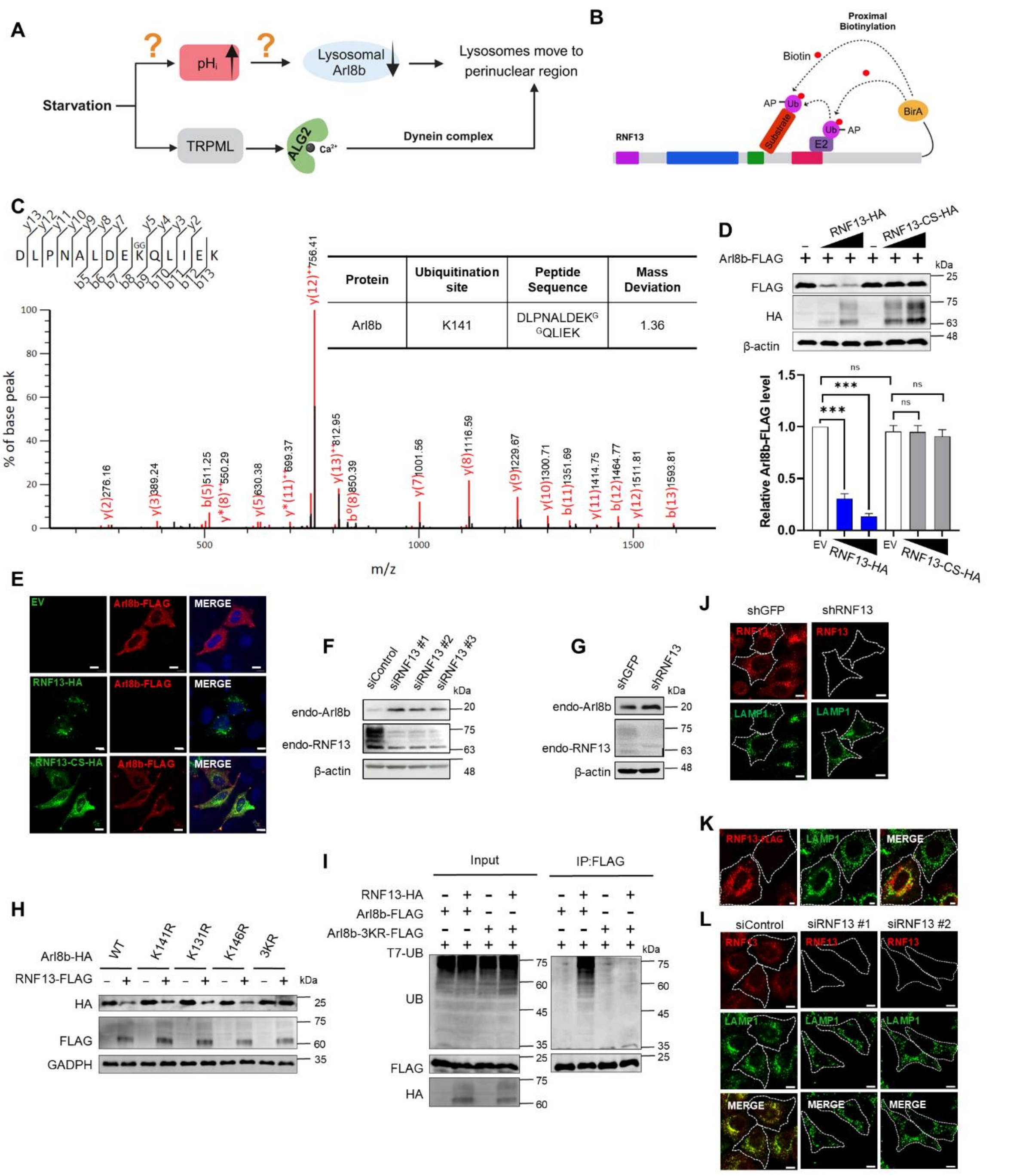


**Figure S1. RNF13 affects lysosomal positioning by promoting ARL8B degradation, related to Figure 1**

(A) Current understating of starvation-induced changes in cytosolic pH ($pH_i$) and the activation of TRPML channels, which facilitate the perinuclear repositioning of lysosomes.

(B) Schematic presentation of the proximity-dependent biotinylation method using BirA-RNF13 fusion protein. Proximal acceptor peptide (AP)-ubiquitin is biotinylated by the biotin ligase BirA fused to RNF13.

(C) MS/MS spectra of a ubiquitin remnant-containing peptide of ARL8B. The sequence of a ubiquitinated peptide derived from ARL8B is indicated, and the fragment ions (b- and y-ions) are labeled. "KGG" represents Gly-Gly-modified lysine (upper panel).

(D) Western blot analysis of indicated proteins in HeLa cell lysates transfected with ARL8B-FLAG in combination with either an empty vector (-) or plasmids encoding RNF13-HA or RNF13 C243S (CS)-HA at increasing concentrations. Relative expression levels are shown as mean ± SD (n = 3). Statistical significance was determined using one-way ANOVA with Dunnett's test. (ns, not significant; ***$p < 0.001$). Protein levels were normalized to β-actin.

(E) Immunofluorescence analysis of ARL8B-FLAG and RNF13-HA in cells. HeLa cells were transfected with ARL8B-FLAG in combination with either an empty vector (-) or plasmids encoding RNF13-HA or RNF13 C243S (CS)-HA. Expression of RNF13-HA dramatically reduces ARL8B-FLAG fluorescence. Scale bar, 10 μm.

(F) Western blot analysis of endogenous ARL8B and RNF13 in HeLa cell extracts transfected with either control siRNA (siControl), or various RNF13 siRNAs.

(G) Western blot analysis of endogenous ARL8B and RNF13 in MCF7 cells stably expressing control shRNA (shGFP), or RNF13 shRNA (shRNF13).

(H) Western blot analysis of ARL8B with K to R substitutions for RN13-mediated degradation. Lysine at 131, 141, or 146 of ARL8B were substituted with arginine. The mutant with K to R substitution at these 3 sites are referred to as 3KR. Lysates of HeLa cells transfected with either ARL8B-HA or ARL8B mutants, along with an empty vector (-) or RNF13-FLAG, were subjected to western blot analysis, indicating that these sites are major sites for ubiquitination.

(I) Impaired the ubiquitination of the 3KR mutant. HeLa cells were transfected with RNF13-HA, T7-Ub, and either ARL8B-FLAG or ARL8B K131/141/146R-FLAG. The cells were treated with MG132 (25 μM) for 6 h and subsequently analyzed for ubiquitination.

(J) Immunofluorescence analysis of LMP1 and RNF13 in MCF7 cells stably expressing control shRNA (shGFP), or RNF13 shRNA (shRNF13). In cells expressing shRNF13, lysosomes are dispersed. Scale bar, 10 μm.

(K) Immunofluorescence analysis of LMP1 and RNF13. HeLa cells were transfected with RNF13-FLAG and immunostained for FLAG and LAMP1 to assess lysosomal distribution. RNF13 overexpression induces the accumulation of lysosomes in the perinuclear region. Scale bar, 5μm.

(L) Immunofluorescence analysis of LMP1 and RNF13. HeLa cells were treated with control siRNA (siControl) or various RNF13-specific siRNAs (siRNF13#1, siRNF13#2) for 72 h, then fixed and immunostained with LAMP1, and RNF13 antibodies. RNF13 knockdown resulted in lysosomal dispersion. Scale bar, 10 μm.

**Supplementary Figure 2.**

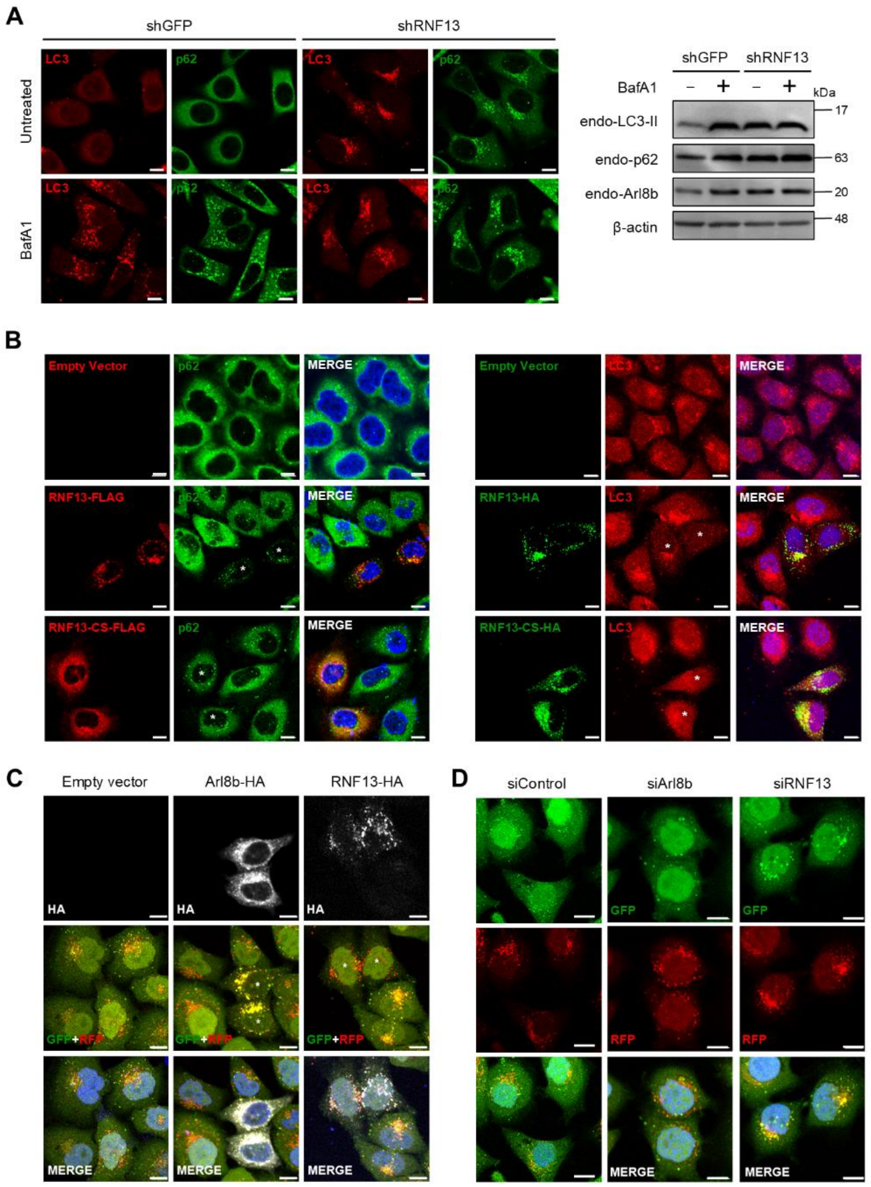


**Figure S2. RNF13-mediated ARL8B degradation stimulates autophagosome-lysosome fusion, related to Figure 1**

(A) Immunofluorescence and western blot analyses of endogenous p62, LC3, and ARL8B following RNF13 knockdown. HeLa cells stably expressing shGFP or shRNF13 were treated with 10 nM Bafilomycin A1 (BafA1) or left untreated. These cells were then either immunostained with LC3 and p62 antibodies (left panel), or immunoblotted with the indicated antibodies (right panel). Scale bar, 5 μm.

(B) Immunofluorescence analysis of autophagy markers, p62, and LC3, following RNF13 overexpression.

HeLa cells were transfected with an empty vector, plasmids encoding RNF13-HA, or RNF13-CS-HA, and then immunostained with the indicated antibodies and DAPI for nucleus staining. Transfected cells with RNF13 constructs are marked with asterisks. Scale bar, 10 μm.

(C, D) Immunofluorescence analysis of mRFP-GFP-LC3. Cells expressing mRFP-GFP-LC3 were either transfected with indicated constructs (C) or subjected to knockdown with indicated siRNAs (D). These cells were then immunostained with an HA antibody (C) and observed for fluorescence signals (C, D). ARL8B overexpression or RNF13 knockdown resulted in prominent yellow signals of mRFP-GFP-LC3, indicating inhibition of autophagosome-lysosome fusion. Scale bars: 10 μm (C) and 5 μm (D).

**Supplementary Figure 3.**

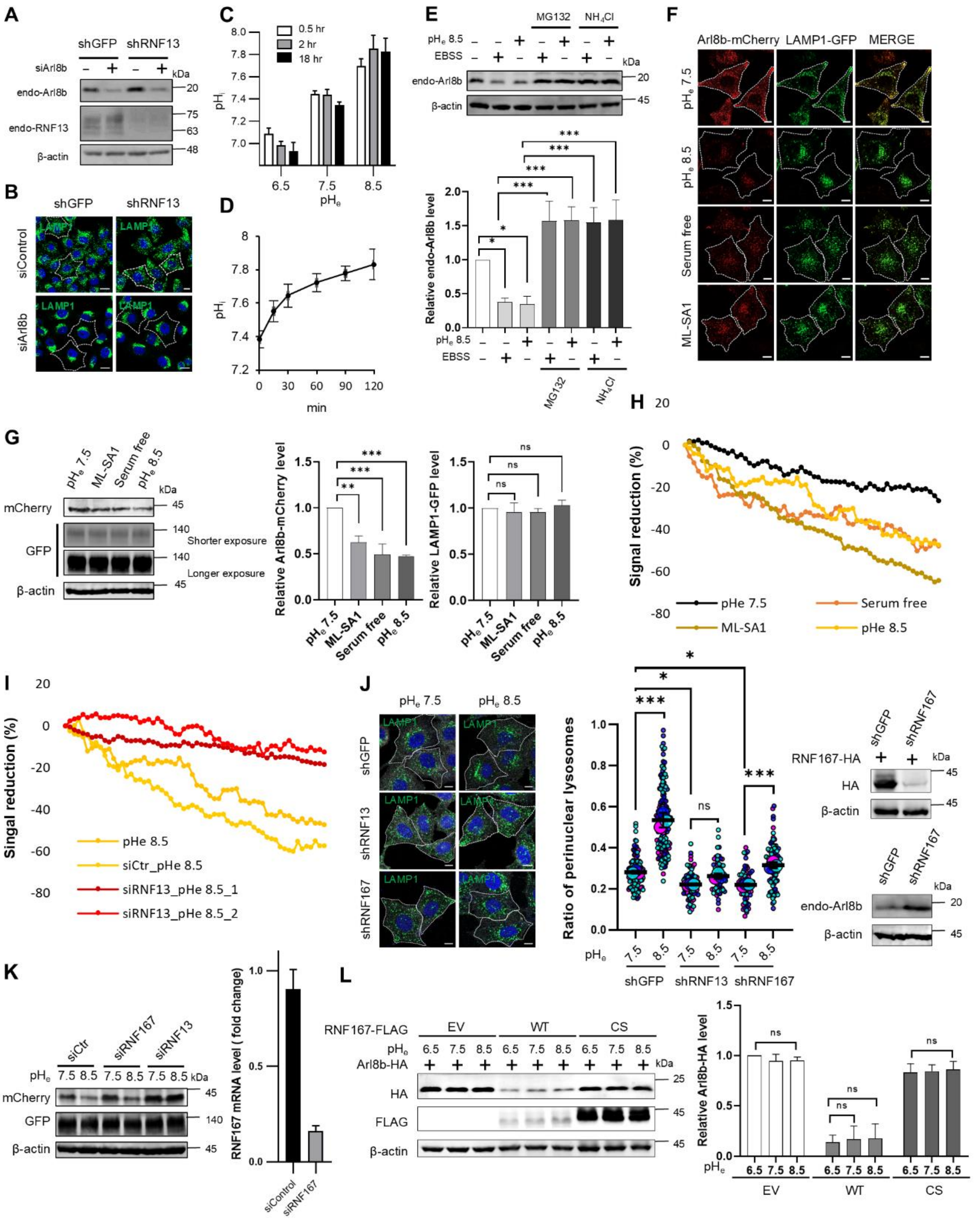


**Figure S3. RNF13 plays a pivotal role in regulating lysosomal positioning, related to Figure 1**

(A) Western blot for endogenous ARL8B and RNF13 and (B) immunofluorescence analyses of LAMP1 in MCF7 cells stably expressing control shRNA (shGFP) or RNF13 shRNA (shRNF13) transfected with control siRNA (siControl) or ARL8B siRNA (siARL8B) for 72 h. Scale bar, 10 μm.

(C, D) Effects of $pH_e$ on $pH_i$. $pH_i$ sensor cells were incubated in media at pH 6.5, pH 7.5, or pH 8.5 for the specified durations (C) or at pH 8.5 for 2 h (D) Fluorescence signals from three replicates (>3 fields each) were converted to $pH_i$ as described in STAR Methods.

(E) Western blot analysis of endogenous ARL8B in HeLa cells treated with either EBSS or pH 8.5, in combination with MG132 or $NH_4Cl$ as specified. Relative expression levels are presented as mean ± SD (n = 3). Statistical significance was assessed using two-way ANOVA followed by Tukey's post-hoc test (*$p < 0.05$; ***$p < 0.001$). Protein expression levels were normalized to β-actin.

(F) Live-cell confocal imaging of ARL8B-mCherry and LAMP1-GFP on HeLa cells stably co-expressing ARL8B-mCherry and LAMP1-GFP following 1 h-treatment under the specified conditions at 37 °C in $CO_2$ incubator. Scale bar, 10 μm.

(G) Western blot analysis of ARL8B-mCherry and LAMP1-GFP in the samples shown in (F). Relative protein levels, normalized to β-actin are presented as mean ± SD from three independent experiments (right panel). Statistical significance was assessed using one-way ANOVA with Dunnett's multiple comparisons test (ns, not significant; **$p < 0.01$; ***$p < 0.001$).

(H) A plot displaying ARL8B-mCherry signals from HeLa cells stably co-expressing ARL8B-mCherry and LAMP1-GFP over a 1-hour treatment period. Conditions include a medium at $pH_e$ 7.5 (Video S1), a medium at $pH_e$ 8.5 (Video S2), a serum free medium (Video S3), and a medium supplemented with 25 μM ML-SA1 (Video S8). Signal quantification is done as described in the Star Methods.

(I) A plot displaying ARL8B-mCherry signals from HeLa cells stably co-expressing ARL8B-mCherry and LAMP1-GFP over 1-hour treatment in medium at $pH_e$ 8.5. Experimental conditions include cells placed in a complete medium ($pH_e$ 8.5: Video S2), cells transfected with control siRNA for 72 h and placed in a medium at $pH_e$ 8.5 (siCtr_$pH_e$ 8.5: Video S4), and cells transfected with RNF13-specific siRNA for 72 h and placed in a medium at $pH_e$ 8.5 (siRNF13_$pH_e$ 8.5_1: Video S5, siRNF13_$pH_e$ 8.5_2: Video S6). Signal quantification is done as described in the Star Methods.

(J) (Left panel) Immunofluorescence analysis of LAMP1 in MCF7 cells stably expressing either shGFP (control shRNA), shRNF13 (RNF13 shRNA), or shRNF167 (RNF167 shRNA) following 2-h incubation in media at $pH_e$ 7.5 or 8.5. Cells were stained with LAMP1 for lysosomes, CD147 for cell boundary and DAPI for nucleus. Scale bar, 10 μm. (Middle panel) A SuperPlot illustrates the ratio of perinuclear LAMP1 to total LAMP1, as detailed in Figure 1E Horizontal lines indicate the mean ± SD of the means from three independent experiments. Statistical significance was evaluated two-way ANOVA with Tukey's multiple comparisons test (*$p < 0.05$; ***$p < 0.001$). (Right upper panel) Western blot analysis of RNF167-HA in MCF7 cells stably expressing shGFP or shRNF167, showing that RNF167 expression is impaired in shRNF167 expressing cells. (Right lower panel) Western blot analysis of endogenous ARL8B in MCF7 cells stably expressing shGFP or shRNF167.

(K) Western blot analysis of ARL8B-mCherry and LAMP1-GFP in HeLa cells stably co-expressing both proteins. Cells were transfected with control siRNA (siCtr), RNF167 siRNA (siRNF167), or RNF13 siRNA (siRNF13) for 72 h and then placed in media at $pH_e$ 7.5 or 8.5. Lysates were collected for subsequent analysis. (Right panel) Knockdown of RNF167 by siRNA was confirmed by qRT-PCR.

(L) Western blot analysis of ARL8B-HA in HeLa cell co-transfected with plasmids expressing ARL8B-HA and either an empty vector (EV), RNF167-FLAG, or RNF167 CS-FLAG. Six hours post-transfection, the cells were incubated in fresh media at pH 6.5, pH 7.5, or pH 8.5 for 18 h and lysates were analyzed. The relative ARL8B-HA levels, normalized to β-actin, were calculated from three independent experiments and presented as mean ± SD. Statistical significance was analyzed by two-way ANOVA with Tukey's test (ns, not significant).

**Supplementary Figure 4.**

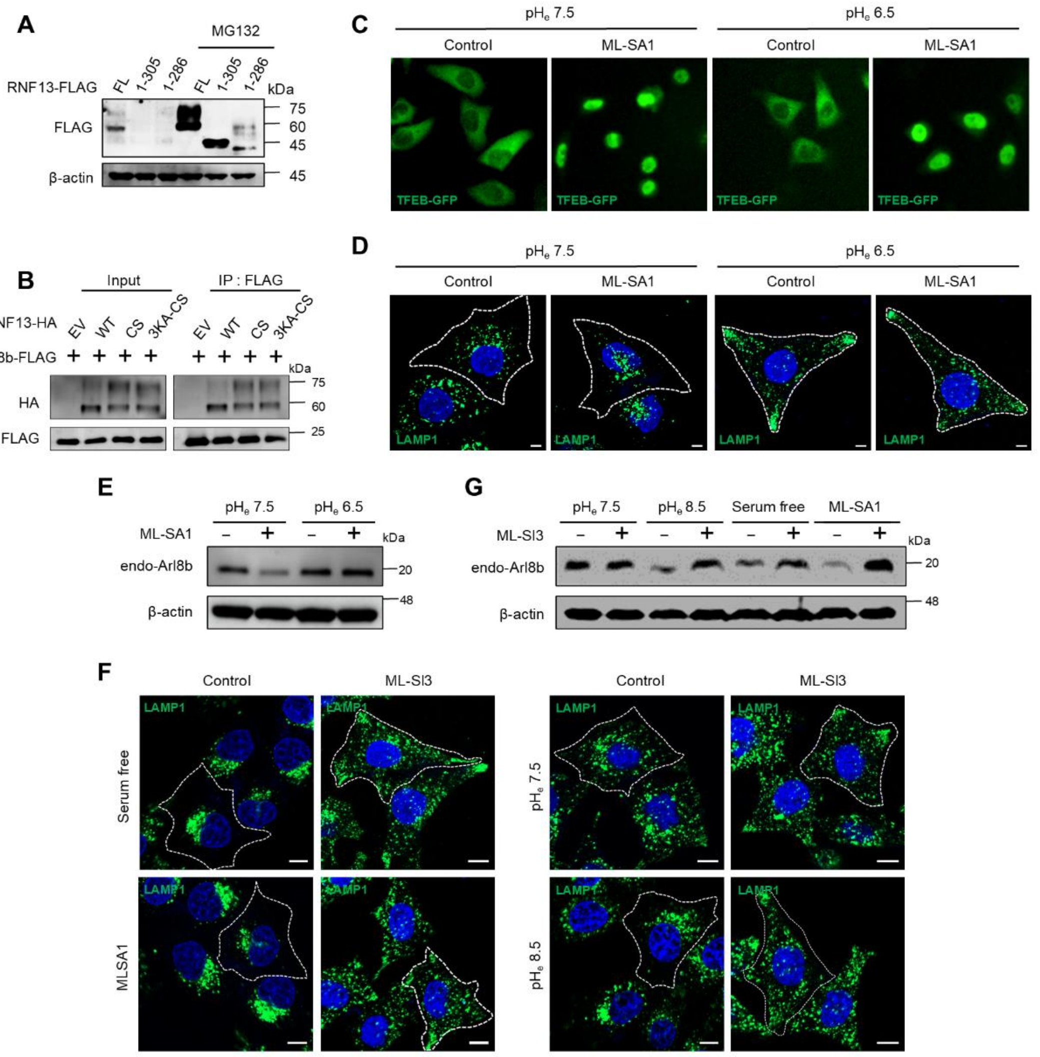


**Figure S4. Effects of different pH$_e$, starvation, ML-SA1, and ML-SI3 on ARL8B levels and lysosome positioning, related to Figures 2 and 4**

(A) Western blot analysis of RNF13-FLAG in HeLa cells expressing RNF13 WT or the indicated mutants, followed by a 6-hour treatment with MG132 (25 μM).

(B) Western blot analysis of RNF13-HA in HeLa cells co-transfected with ARL8B-FLAG, along with indicated RNF13 constructs for 24 h, treated with MG132 (25 μM) for 6 h prior to harvesting, and analyzed by IP with FLAG antibody.

(C) Immunofluorescence analysis of TFEB-GEP in media at different pH, showing that cytosolic acidification does not inhibit the translocation of TFEB to the nucleus upon ML-SA1 treatment. HeLa cells were transfected with a TFEB-GFP construct and treated with DMSO or ML-SA1 (25 μM) in complete media at either pH 7.5 or pH 6.5 for 2 h.

(D, E) Immunofluorescence analysis of lysosome positioning (D) and western blot of endogenous ARL8B (E) upon ML-SA1 treatment under acidic conditions. HeLa cells were treated with DMSO or ML-SA1 (25 μM) in complete media at pH 7.5, or pH 6.5 for 2 h. The cells were then immunostained with LAMP1 (D), or analyzed by immunoblotting (E). Scale bar, 5 μm. .

(F, G) Immunofluorescence and western blot analyses of TRPML inhibition under alkalinization or starvation on lysosomal positioning and ARL8B degradation. HeLa cells were pretreated with DMSO (control) or ML-SI3 (25 μM) for 30 min. The media was then replaced with serum-free media, or media containing ML-SA1 (25 μM), or fresh media at pH 7.5 or pH 8.5 for 2 h, in the presence or absence of ML-SI3. After treatment, the cells were either by immunostained with a LAMP1 antibody and DAPI for nuclear staining (F) or analyzed by Western blot of endogenous ARL8B (G). Scale bar, 10 μm.

**Supplementary Figure 5.**

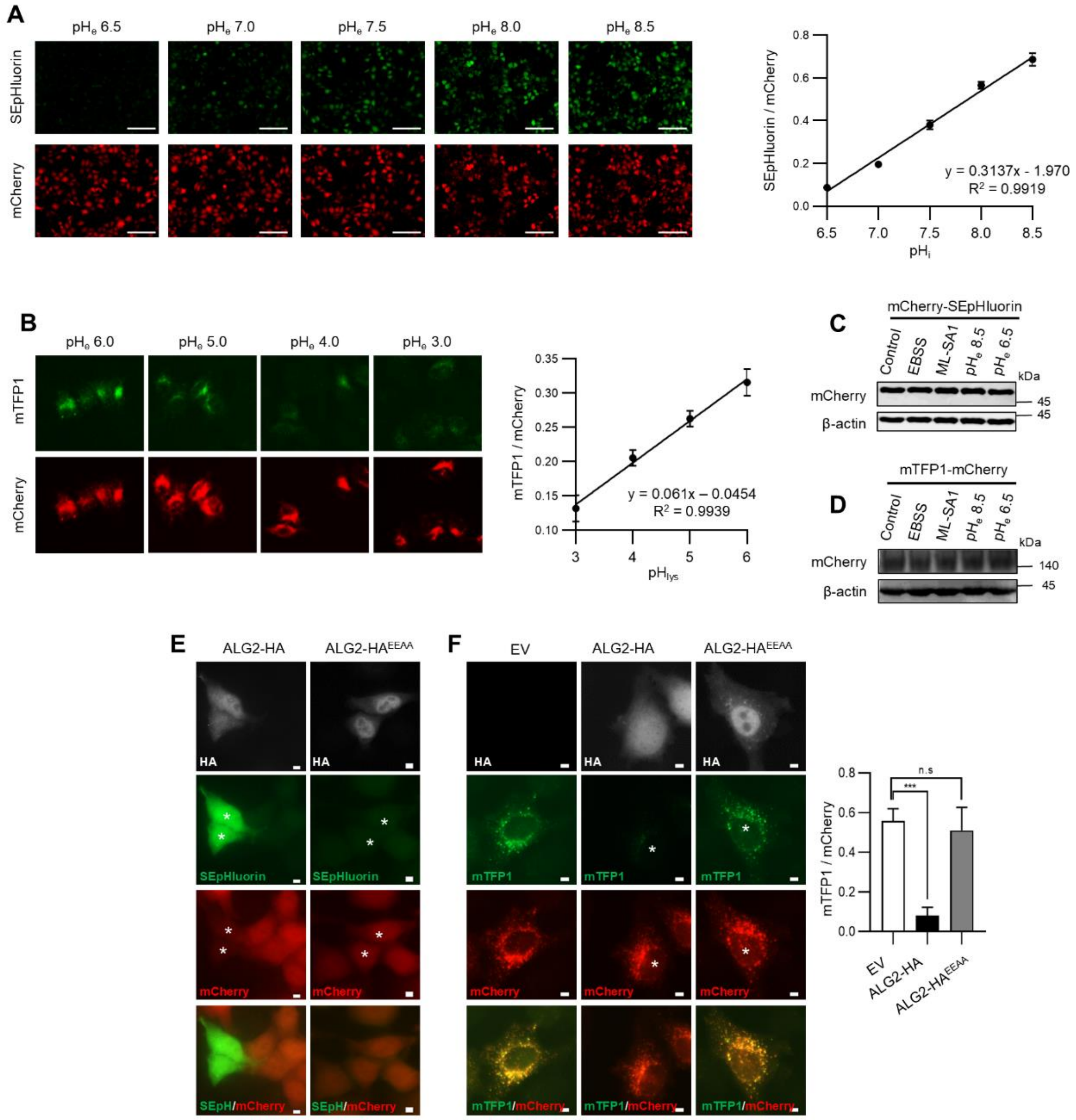


**Figure S5. The role of ALG2 in regulation of $pH_i$ and $pH_{lys}$, related to Figures 5-7**

(A, B) (Left panel) Representative images of $pH_i$ sensing HeLa cells stably expressing mCherry-SEpHluorin (A) or $pH_{lys}$ sensing HeLa cells stably expressing mTFP1–hLAMP1–mCherry (B) at different pHs. Scale bar, 200 µm. (Right panel) Standard curves were generated by incubating $pH_i$ sensing cells (A) in buffers at pH 6.5 - 8.5, or $pH_{lys}$ sensing (B) in buffers at pH 3.0 - 6.0. Data are presented as mean ± SD of SEpHluorin/mCherry ratio from at least three fields per pH point (A), or mTFP1/mCherry ratio from 20 cells per pH point with three independent replicates (B).

(C, D) Western blot analysis of pH sensor protein levels in HeLa cells stably expressing mCherry-SEpHluorin (C) or mTFP1–hLAMP1–mCherry (D) after treatment under the specified conditions for 2 h. The levels of $pH_i$ and $pH_{lys}$ sensor proteins were not affected by various treatments employed.

(E, F) Immunofluorescence analysis of the sensor proteins in HeLa cells stably expressing mCherry-SEpHluorin (E), or mTFP1–hLAMP1–mCherry (F) transfected with ALG2-HA or a $Ca^{2+}$ binding-deficient ALG2 mutant, ALG-2$^{E47A/E114A}$-HA. Asterisks mark transfected cells. Overexpression of ALG2-HA but not the mutant dramatically increases SEpHluorin signals, indicating the elevated $pH_i$ (E). In panel F, overexpression of ALG2-HA but not the mutant reduced $pH_{lys}$. The ratios of green to red signals were obtained from five cells per condition. Statistical significance was determined using one-way ANOVA with Dunnett's for multiple comparisons test (ns, not significant; ***$p < 0.001$).

**Supplementary Figure 6.**

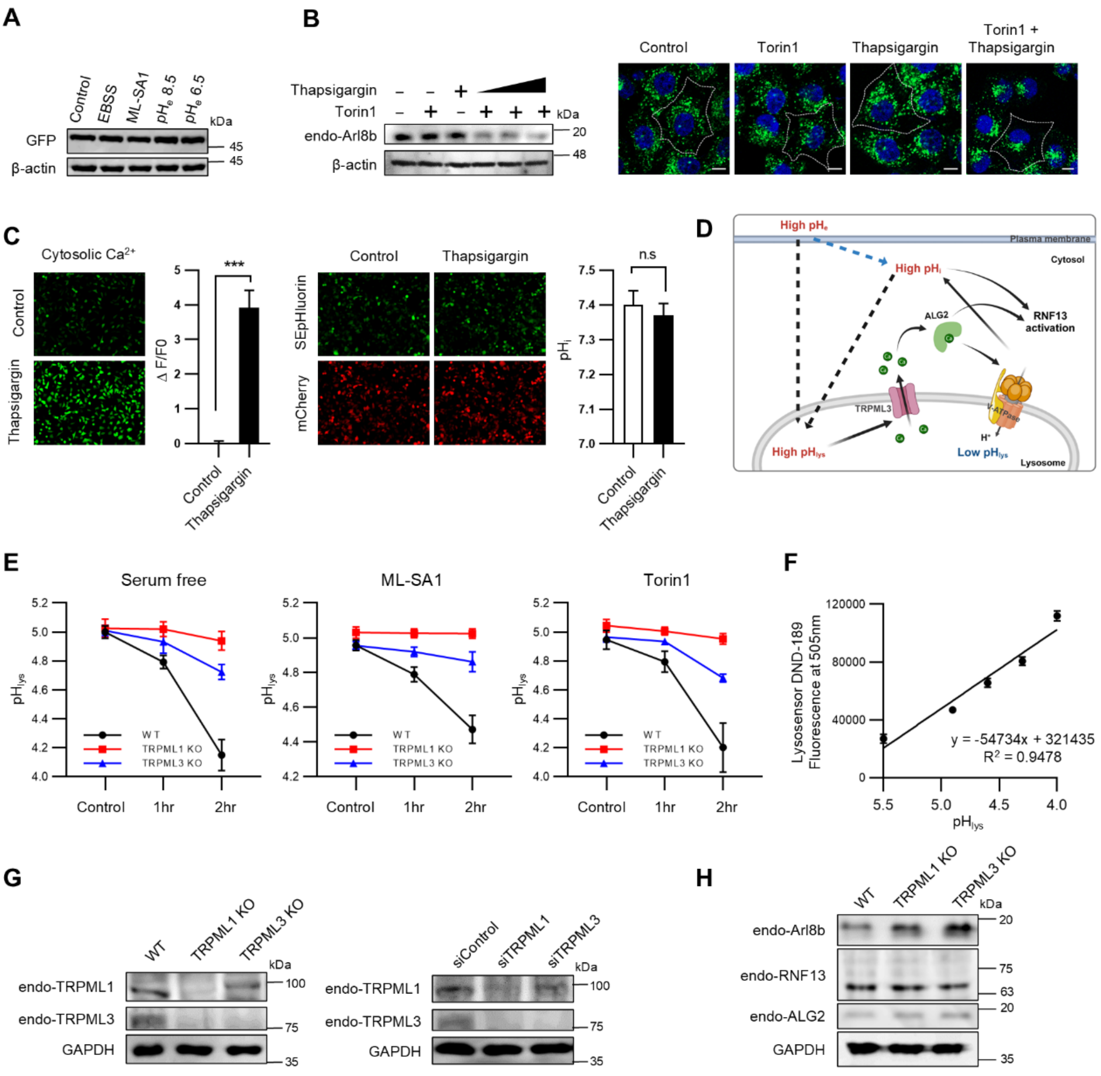


**Figure S6. Cytosolic alkalinization and increased cytosolic $Ca^{2+}$ are essential for ARL8B degradation and perinuclear positioning of lysosomes, and the involvement of TRPML channels in lysosomal acidification under treatments, related to Figures 6 and 7**

(A) Western blot analysis of the $Ca^{2+}$ sensor GCaMP6f-GFP protein levels in HeLa cells stably expressing GCaMP6f-GFP after 2-h treatment under the specified conditions. The GCaMP6f levels remained unchanged 2 h post treatment across various conditions.

(B) Western blot and immunofluorescence analyses of ARL8B levels and lysosomal positioning following co-treatment with Torin 1 and thapsigargin. HeLa cells were treated with DMSO (control), Torin1 (400 nM), thapsigargin (2 μM), or Torin1 (400 nM) in combination with thapsigargin rainging from 1 to 4 μM for 2 h. Cells were used for either western blot analysis (left panel) or immunostained with a LAMP1 antibody and DAPI for nuclear staining (right panel). Scale bar, 10 μm.

(C) Thapsigargin increases cytosolic $Ca^{2+}$ levels without affecting $pH_i$. HeLa cells stably expressing GCaMP6f (left panel) or mCherry-SEpHluorin (right panel) were treated with thapsigargin (2 μM) for 2 h. Scale bar, 200 μm. Signals from three replicates, each with at least three fields, were analyzed using an unpaired Student t-test (***$p < 0.0001$: ns: not significant).

(D) A model illustrating how alkaline $pH_e$ triggers an immediate increase followed by a subsequent decrease in $pH_{lys}$. It also depicts that alkaline $pH_e$-induced ALG-2 activation and $pH_i$ increase enhances RNF13 activity. The image was created using BioRender.com.

(E) Effects of specified conditions on $pH_{lys}$ in HAP1 (WT, TRPML1 KO, and TRPML3 KO) cells. Cells were treated with serum-free media, ML-SA1 (25 μM), or Torin1 (400 nM) for the indicated times. Cells were treated with 2 μM Lysosensor DND-189 for 1 h. Signals from at least three fields per condition were quantified, with three replicates per condition.

(F) Standard curve for Lysosensor-based pH measurements.

(G) Western blot analysis of endogenous TRPML1 and TRPML3 levels from HAP1 (WT, TRPML1 KO, and TRPML3 KO) cells (left panel) or HeLa cells transfected with control siRNA, TRPML1-siRNA, or TRPML3-siRNA for 72 h (right panel).

(H) Western blot analysis of endogenous proteins from HAP1 (WT, TRPML1 KO, and TRPML3 KO) cell lysates, immunoblotted with the indicated antibodies.

### Supplementary Figure 7.

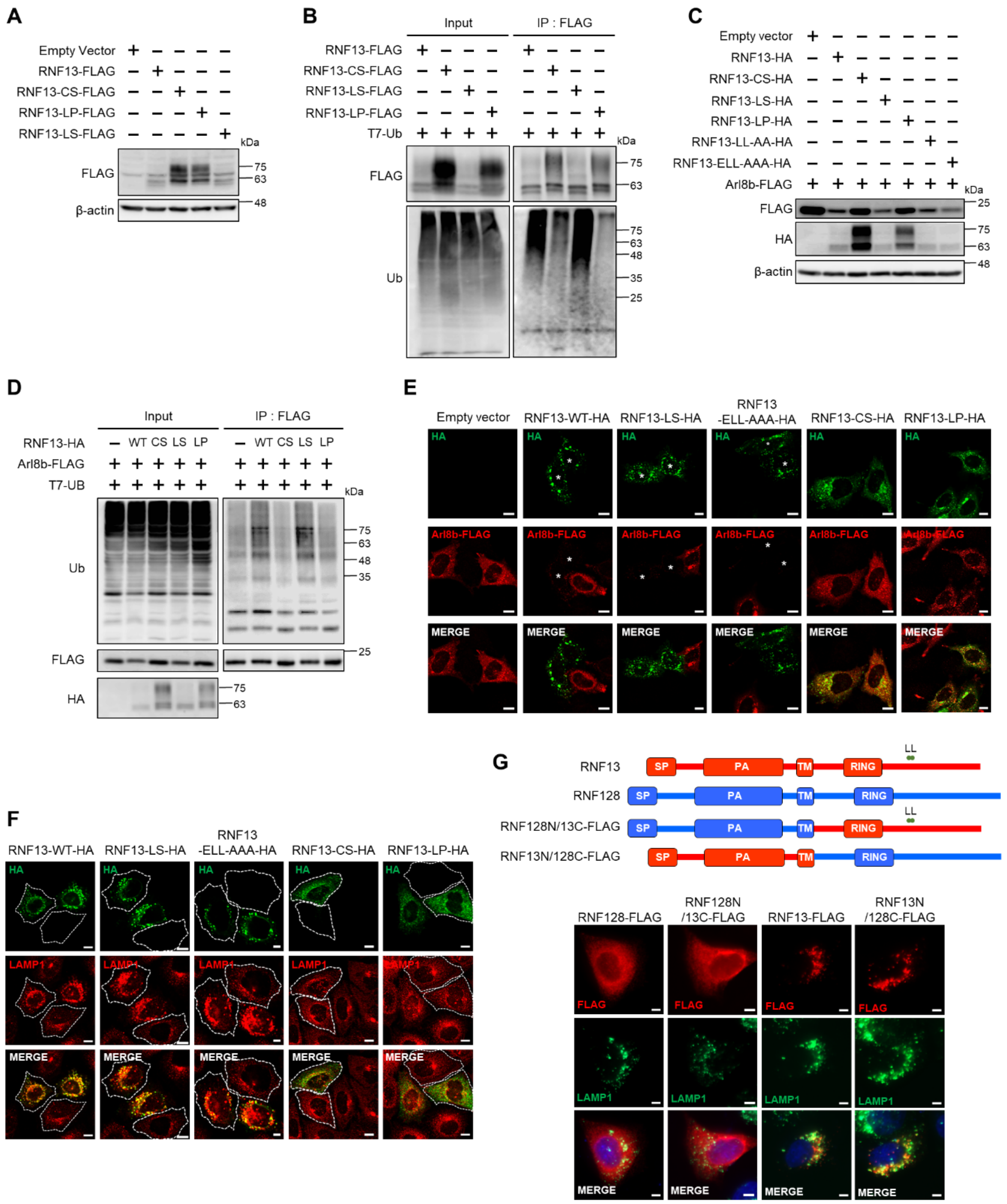


**Figure S7. The loss of ubiquitin ligase activity in RNF13 L312P is implicated in the pathogenesis of DEE-73, related to Figure 2**

(A, B) Western blot and ubiquitination analyses revealed a loss of ligase activity of RNF13 L312P mutant. HeLa cells transfected with indicated constructs and analyzed by immunoblotting with FLAG antibodies

(A) and ubiquitination assay as described in Figure 1C (B). RNF13 CS, LP, and LS refers to RNF13 C243S, L312P, and L311S mutants, respectively.

(C, D) Western blot analysis and ubiquitination assay of HeLa cells transfected with the indicated constructs. HeLa cells were co-transfected with ARL8B-FLAG and the indicated constructs, then analyzed by immunoblotting (C) or ubiquitination assay (D).

(E) Immunofluorescence analysis of HeLa cells co-transfected with ARL8B-FLAG and RNF13 mutants. Scale bar, 10 μm.

(F) Immunofluorescence analysis of lysosomal positioning of HeLa cells transfected with indicated RNF13 mutants. Scale bar, 10 μm.

(G) RNF13 and RNF13N/128C are localized to lysosomes. Schematic presentation of RNF13 and RNF128 fusion constructs (upper panel). SP; signal peptide, PA; protease-associated domain, TM; transmembrane domain, RING; a really interesting new gene domain.

HeLa cells were transfected with the indicated constructs for 24 h, fixed, and then immunostained with LAMP1 antibodies (lower panel). Scale bar, 10 μm.
